# Motor cortical output for skilled forelimb movement is selectively distributed across projection neuron classes

**DOI:** 10.1101/772517

**Authors:** Junchol Park, James W. Phillips, Jian-Zhong Guo, Kathleen A. Martin, Adam W. Hantman, Joshua T. Dudman

**Author notes:** co-first authors (Junchol Park, James W. Phillips, Jian-Zhong Guo). co-senior authors (Adam W. Hantman, Joshua T. Dudman).

## Abstract

The interaction of descending neocortical outputs and subcortical premotor circuits is critical for shaping the skilled movements. Two broad classes of motor cortical output projection neurons provide input to many subcortical motor areas: pyramidal tract neurons (PT), which project throughout the neuraxis; and intratelencephalic neurons (IT), which project within cortex and subcortical striatum. It is unclear whether these classes are functionally in series or whether separable components of descending motor control signals are distributed across these distinct classes of projection neurons. Here we combine large-scale neural recordings across all layers of motor cortex with cell-type specific perturbations to study cortically-dependent mouse motor behaviors: kinematically-variable manipulation of a joystick and a kinematically-precise reach-to-grasp. We find that striatum-projecting IT neuron activity preferentially represents amplitude whereas pons-projecting PT neurons preferentially represent the variable direction of forelimb movements. Thus, separable components of descending motor cortical commands are distributed across motor cortical projection cell classes.

**One-sentence summary:** Separable components of cortical motor commands are distributed across distinct glutamatergic projection neuron cell-types.

## Introduction

In mammals, descending motor control signals from the neocortex are carried via several classes of molecularly-defined output projections neurons (*1, 2*). Pyramidal tract (PT) neurons project directly to the midbrain, brainstem, and spinal cord, along with other descending systems (*2–5*). Intratelencephalic (IT) projection neurons of layer 5 and layer 2/3 (*2, 3, 6–8*) project within the forebrain and prominently target subcortical striatum in both hemispheres (often referred to as ‘corticostriatal’ (*6*)). PT neurons are a primary output cell type throughout vertebrate lineage (*1*), whereas expansion and diversification of IT neuron populations appears to be a major contributor to changes in motor cortical cell types in mammals (*9*).

Cerebellum and basal ganglia are subcortical targets of motor cortical output that are thought to be particularly critical for the fine control of skilled limb movements (*10–13*). Cerebellum receives cortical input indirectly via ascending mossy fibers from brainstem nuclei including the basal pons. Basal pons receive dense input from nearly all PT neurons from the primary motor cortex (and other cortical areas) (*14–16*). Inactivation of basal pons produces fine targeting deficits in forelimb movements (endpoint errors) while leaving gross kinematics (speed and amplitude) relatively unaffected (*17*). Cerebellum is thought to be the brain locus in which forward models are computed - transforming copies of motor commands into predictions used for feedback control (*18–20*). Disruption of cerebellar function leads to deficits in the appropriate targeting of movement direction (*20*) supporting such an interpretation.

Basal ganglia circuits are critical for controlling the execution of skilled forelimb movements (*21–26*) and are closely associated with regulation of the amplitude and speed of movement (*26–28*). Striatum, the forebrain input nucleus of basal ganglia (*7*), is unique amongst subcortical motor areas in being a target of both IT and PT neurons (*3, 6, 7, 29*). A number of models have been proposed to account for the computations in basal ganglia that underlie its role in specifying the speed and amplitude of movement (*10, 27, 28, 30–32*). A common feature of these models is that descending cortical motor commands are carried to striatum where basal ganglia circuits may modulate the gain (*27*) of descending motor commands (termed movement vigor (*26, 33*)) or implement a closed-loop feedback to shape movement kinematics (*31*) and/or act as a primary source of motor commands for stereotyped movement trajectories (*28, 32*). In contrast to basal pons deficits, inactivation of dorsal striatum modifies movement speed and amplitude while often leaving movement target direction unaffected (*21, 22, 24, 34*).

Thus, key computations proposed for cerebellar and basal ganglia circuits depend upon copies of descending cortical motor commands carried by motor cortical projection neurons. On the one hand it has generally been believed that PT neurons of motor cortex may be the pathway in which cortical motor commands emerge and are conveyed to subcortical targets to mediate diverse aspects of motor control (*4, 10*). From this perspective, it has been proposed that IT neurons may also exert influence on movement via PT output pathways; for example, it has been argued that ‘pre-movement’ (motor planning) activity in IT neurons is transformed into motor commands in PT populations (*35*). This is consistent with asymmetric connectivity in the motor cortex that exhibits a strong IT→PT bias (*36*) and the fact that PT neurons elaborate collaterals within many subcortical targets (*2–4*). From this perspective IT inactivation is expected to result in similar or smaller consequences on movement as compared to PT inactivation (*35*). However, many studies have reported substantial movement execution-related activity in non-PT cell types in the motor cortex (*8, 37–42*). Moreover, it is less clear from this perspective why inactivation of two different subcortical targets, striatum and pons, result in different (and often dissociable) effects on forelimb movement execution if the majority of motor command information arises from a largely shared population of PT neuron inputs. For these reasons we entertain a modified perspective that can potentially reconcile these data.

Rather than PT neurons being the primary or even sole locus at which cortical activity is transformed into descending neocortical motor commands, it is possible that descending motor commands are distributed across corticostriatal IT neurons and corticopontine PT neurons. From the perspective of downstream perturbations and anatomical differences, one putative division would be IT neuron populations carry information about movement amplitude whereas corticopontine PT neuron populations are preferentially involved in control of movement direction. This may be consistent with movement amplitude encoding in striatum requiring information related to the intensity of muscle activation. In contrast, forward model computations in cerebellum may depend more critically upon information about which muscle groups are activated (e.g. flexor/extensor ratio which determines movement direction) - information known to be present in PT populations (*4*). This revised perspective is potentially distinguishable from a model in which intracortical IT→PT projections transform premotor activity into motor commands because it (1) implies differential encoding of movement kinematic parameters in IT and PT populations; and (2) predicts dissociable consequences on movement execution during inactivation of each projection cell class. To date, the putative differential encoding of movement parameters and dissociable effects of IT and PT inactivation on movement execution have been little studied.

Here we sought to address this question by combining large-scale neural recording across the entire motor cortical depth and striatum with cell-type specific identification and perturbation in the context of mice performing skilled forelimb motor tasks that were either highly variable or highly consistent in movement direction and amplitude. This allowed us to explore the information about movement kinematics observed in molecularly distinct corticostriatal IT (Tlx3+) motor projection neurons as compared with corticopontine PT (Sim1+) neurons. We found that IT neuron activity was a rich source of information preferentially about movement amplitude and as compared to corticopontine PT neurons which were relatively more informative about movement direction than amplitude. These neural correlates were consistent with partially dissociable effects of cell-type specific inactivation. Tlx3+ IT neuron inactivation produced a large attenuation of movement speed and amplitude whereas inactivation of corticopontine PT neurons produced alterations in movement direction with relative minor changes in amplitude. These data provide evidence for a multimodal efference system in primary motor cortex in which separable components of descending motor control signals for the same effector are distributed across molecularly distinct projection neuron classes.

## Results

To assess the potential differential encoding and function of motor cortical projection neuron classes we studied a task in which mice make skilled, but highly variable forelimb movement of a joystick to collect delayed reward. Mice were trained to make self-initiated (uncued) movements past a threshold (either directed away from or towards the body) in order to obtain a delayed (1 s) water reward (Fig. 1a-b; Supplementary Video 1). Changing the required movement threshold across blocks (near-far-near) led mice to adjust reach amplitude across blocks (Fig. 1c, repeated measures ANOVA, F_2,16_ = 13.28, p=4.0×10^-4^, between blocks Kruskal-Wallis (KW) test, p=0.01; n=6 mice, 10 sessions) and elicited a broad distribution of movement amplitudes (Fig. 1d; mean: 8.1 ± 4.4 s.d. mm, max: 24.2 mm). Reward could be elicited by suprathreshold joystick movements in any direction in the 2d plane of the joystick. Mice preferentially used movements with variable direction although biased towards movements along (towards or away from) the body axis (Fig. 1e).

**Fig. 1.**
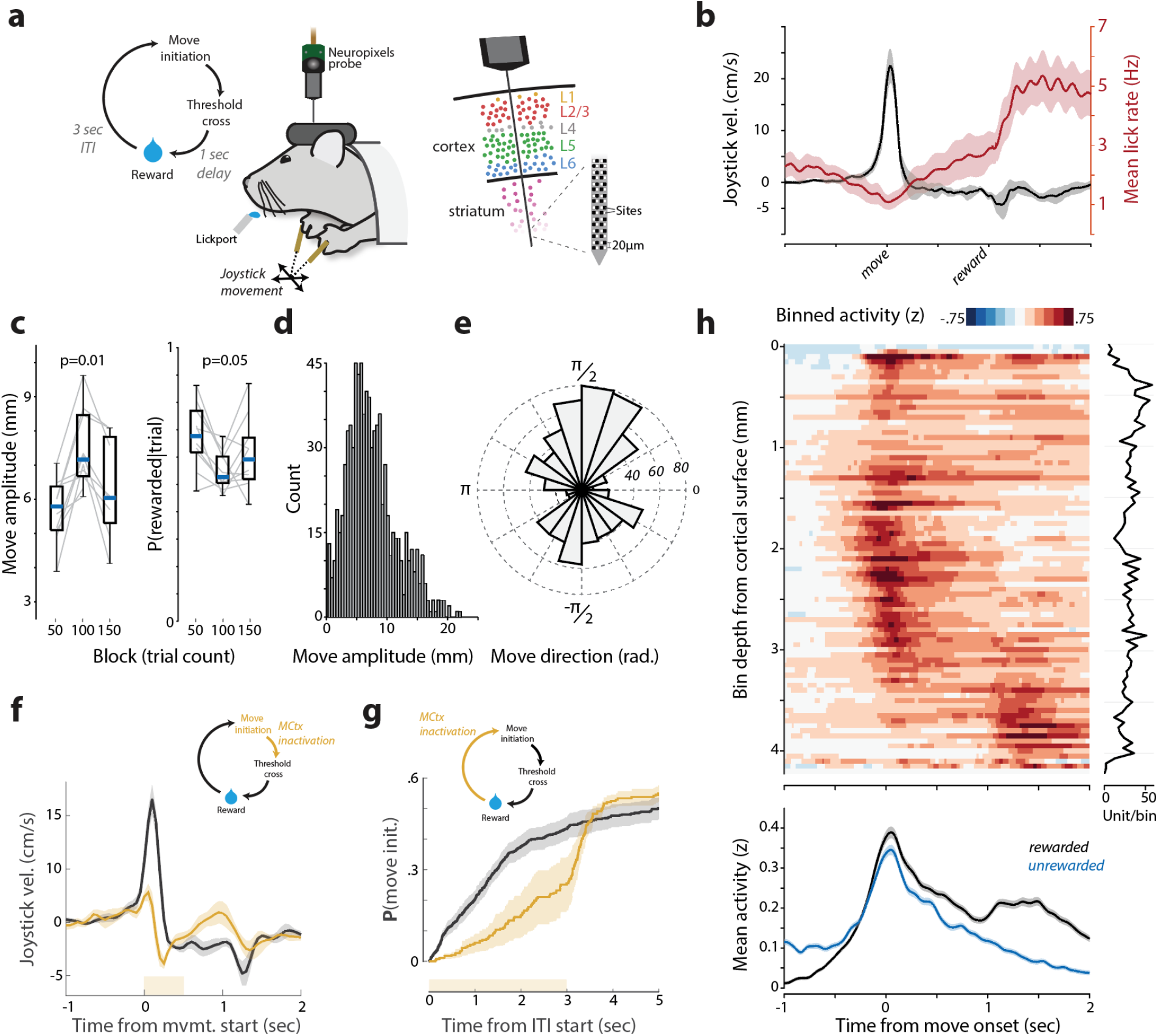
Distributed task-related neural activity in a variable amplitude operant task. **a**, Schematic of behavioral task and Neuropixels probe recordings from mouse MCtx^FL^. **b**, Outward joystick velocity and lick rate aligned to threshold crossing for 10 sessions (6 mice). Shaded area indicates the s.e.m. **c**, Movement amplitude as a function of threshold. Probability of initiating a movement of the correct amplitude. **d-e**, Distribution of movement amplitude and direction. **f**, Closed-loop inactivation of MCtx^FL^ in VGAT-ChR2 mice (500 ms duration, yellow bar) triggered on movement initiation. Joystick velocity on control (black) and inactivated (yellow) trials. **g,** Cumulative probability of movement initiation for control (black) and open-loop inactivation (yellow) trials. N=3 mice, 2 sessions/mouse. **h**, Spike density functions of neural activity aligned to movement onset and binned (50 µm bins) according to recording depth. Plot at right shows the number of units/bin. Task-related neural activity was widely distributed across the depth of recordings. Lower plot shows the mean activity across all units for movements that were rewarded (black) vs. comparable magnitude movements (matched median) that were unrewarded (cyan) and lacked the later reward consumption related modulation.

This task, and the ability to adapt movement amplitude to a changing threshold across blocks, depends upon basal ganglia function in mice (*22, 43*); however, motor cortical dependence was unclear. Thus, we first asked whether activity in the forelimb motor cortex (MCtx^FL^) was critical for performance by using optogenetic inactivation (*44*) during or prior to movement execution (*22*). First, we allowed the initiation of movement to occur and then rapidly triggered (*22*) optical inactivation of MCtx^FL^ using VGAT-ChR2 mouse (*45*). As in other mouse forelimb operant tasks, *e.g.* joystick (*46*) or reach-to-grasp (*44, 47*), we found that MCtx^FL^ was critical for normal movement execution (Fig. 1f, Supplementary Video 1; ANOVA, amplitude; F_1,10_ = 11.33, p = 0.007; speed; F_1,10_ = 47.55, p = 4.22×10^-5^). In addition, tonic inactivation of MCtx^FL^ prior to movement initiation significantly reduced the probability of reach initiation (Fig. 1g, ANOVA, F_1,10_ = 8.38, p = 0.016). Finally, optogenetic activation of combined layer 5 MCtx^FL^ output pathways (using the Rbp4-cre mouse line (*2, 48*)) was sufficient to increase the speed and amplitude of forelimb movements (Fig. S1).

To examine neural activity across all layers of MCtx^FL^ during forelimb movements we used Neuropixels probes (*49*). A total of 2416 well-isolated and histologically-verified (see Methods) single units were recorded across MCtx^FL^ (N=1212) and underlying striatum (STR; N=1204 units, Fig. 1a, fig. S2; 220 ± 53 s.d. units per session, total 11 recording sessions). This task allowed us to isolate in time neural activity related to outward forelimb movements from modulation of activity during delayed reward collection (Fig. 1h). Task-related activity was distributed across the entire recording depths including many single units in MCtx^FL^ and dorsal STR (dSTR) with robust movement-timed activity (Fig. 1h, figs. S3 & S4). Activity modulated during reward collection could be revealed by comparing unsuccessful movements (during the ITI or that failed to hit the amplitude threshold) to those that did yield a reward (Fig. 1h lower). Population activity primarily differed around the time of reward collection and was relatively unchanged during execution of approximately matched amplitude movements. Consistent with this difference, when aligned on reward delivery a substantial number of units in MCtx^FL^ were robustly modulated (Fig. S3a).

### Heterogeneous distribution of motor command activity across recording depths

We next sought to examine how activity related to forelimb movement kinematics was distributed anatomically along our electrode recording tracks. First, electrode tracks of individual recording sessions were visualized and anatomical positions were registered to a standard brain atlas (Fig. 2a, fig. S2a-b; see Methods). Next, we confirmed that the population of neurons along our recording track contained robust information about movement kinematics even with these highly variable (trial to trial) movement trajectories. We developed an approach to train linear decoders of movement kinematics assessed on held out trials (see methods). Linear decoders were able to capture much of the variance in observed joystick trajectories even for single trials (Fig. 2b-c). Notably, we observed that decoding performance appeared good independent of the direction of movement (which trials were held out). This decoding performance suggests the presence of information about both movement amplitude and direction in neural population activity. To examine the independent encoding of direction and amplitude we developed a modified approach (see Methods) to identify two targeted, orthogonal dimensions of population activity that best captured activity modulation correlated with movement amplitude (termed ‘AMP’ dimension) or direction (‘DIR’). In all datasets (N=11; N=7 mice) we found robust neural tuning to amplitude and direction when projected along independent AMP and DIR modes of neural population activity (Fig. 2d-e).

**Fig. 2.**
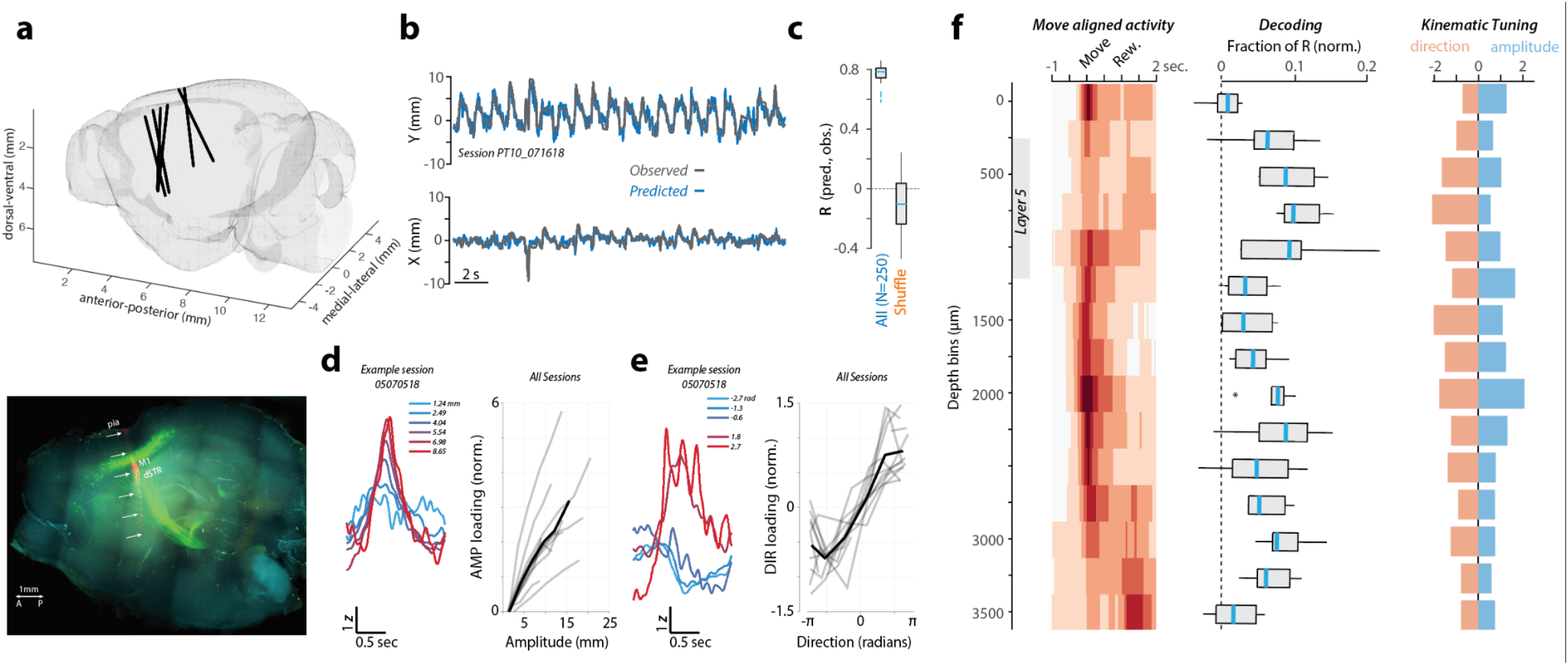
Inhomogeneous representation of movement kinematics across corticostriatal de. **a**, *Upper*, Probe tracks reconstructed in standard brain reference coordinates (see methods). *Lower*, Labeling of pons-projecting PT neurons (green) and the probe tract (red)imaged in a cleared hemibrain (see figs. S2 & S3). Scale bar=1mm. **b**, Decoded (blue) vs observed (gray) movement profiles for 20 rewarded (concatenated, permuted order) joystick movements (see methods). **c**, Cross validation decoding performance compared to shuffle control (Pearson correlation; see Methods). **d-e**, Targeted dimensionality reduction (see Methods) identified two orthogonal dimensions of population activity that encoded amplitude (d) and direction (e). For each: *Left*, example session mean projection of movement aligned activity on AMP (d) or DIR (e) dimension as function of movement amplitude (d) or direction (e; mean value for quintile shown in color legend) quintiles. *Right*, Integrated perimovement modulation of activity (loading) along AMP (d) and DIR (e) dimensions as a function of the average amplitude (d) and direction (e) of movement (offset normalized). Individual sessions: grey; Mean: black. **f**, Neural correlates as a function of depth. For each depth bin (250µm): *Left*, movement-aligned activity; *Middle*, relative contribution to decoder performance (see Methods); *Right*, tuning of AMP and DIR activity dimensions to movement amplitude and direction quintiles.

The use of linear methods in the full dimensionality of population activity (empowered by simultaneous recording of large populations), allowed us to assess how individual units contributed to decoding of movement trajectories and tuning to amplitude and direction as a function of anatomical position. We next compared 3 different aspects of the recorded data in 250 µm bins of depth from the cortical surface for all recording sessions. For each group of units per depth bin we compare the mean peri-movement activity time histogram (Fig, 2f *left*, ‘Move aligned activity’; same as Fig. 1h but with broader spatial bins), the contribution to decoder performance (fraction of explained variance; ‘Decoding’), and the slope relating activity along AMP or DIR dimensions with the amplitude or direction of movement (‘Kinematic tuning’; Fig. 2d-e; see Methods).

Across the population of recorded neurons, the largest decoding contributions were found in the region of Layer 5 of MCtx^FL^ and dorsal STR (Fig. 2f, *middle*) consistent with previous data showing correlates to movement kinematics in dSTR in this task (*22, 34*). In contrast, a relative dearth of movement-related activity and modest decoder performance contribution was observed in ventral striatum. Whereas decoding and tuning to movement kinematic features was, in general, distributed broadly across depth, there was bias toward an increased tuning to movement direction relative to amplitude in intermediate to deep Layer 5 of motor cortex (Fig. 2f *right*, orange bars). To further confirm laminar inhomogeneity of movement-related activity, we also computed the principal component of cortical population activity and found that movement-related modulation of activity was preferentially loaded in upper layers of motor cortex (superficial layer 5 and up) (Fig. S5).

### Molecularly and anatomically defined projection cell classes during recording

IT and PT neurons have partially overlapping, but characteristic laminar positions in neocortex (*1–3, 6*) suggesting that the laminar inhomogeneity in the encoding of movement kinematics (Fig. 2f) could reflect differences in the neural correlates in IT and PT projection cell classes in layer 5. Thus, we next used optogenetic tagging (*35, 50, 51*) to distinguish striatum-projecting layer 5a IT neurons from pons-projecting deep layer 5b PT neurons while simultaneously recording population activity across depths to allow identification of AMP and DIR encoding dimensions. Mouse lines exploiting cell-type specific expression of Tlx3 and Sim1 (*48*) allow molecular access to distinct layer 5 IT and PT subtypes, respectively (*2, 35*). PT projection neurons are a diverse class that project to partially distinct subsets of downstream regions (*2*). Thus, to achieve labeling of pons-projecting PT neurons we used a retrograde virus (rAAV2-retro (*52*)) with conditional expression of the inhibitory opsin FLInChR (*53*) injected into the brainstem (pons) of Sim1-cre mice (*48*). We used a robust and rapid optogenetic inhibitor (to mitigate against confounds due to extensive cortical recurrent excitation (*50*)) that produces efficient inactivation of PT neurons (*53*) to identify neurons in awake animals.

This strategy resulted in robust expression of an inhibitory opsin in brainstem-projecting PT neurons located within layer 5 of MCtx^FL^ (Fig. 2a, fig. S6). A total of 111 units were identified as ‘tagged’ (PT^+^). In PT^+^ subset, activity was inhibited at short latency with half-maximal inhibition occurring within 15 ms on average (median latency < 10 ms) after illumination onset, and the inhibition lasted below the half-maximal level for on average 931 ms of the 1-sec laser (Fig. 3a, c, d, fig. S7; see Methods for statistical criteria for tagging). To identify layer 5 IT neurons we used a similar retrograde labeling strategy with FLInChR injected into the dorsal striatum of Tlx3-cre (*48*) mice. This led to expression of FLInChR in striatum-projecting IT neurons within layer 5 of MCtx^FL^ (Fig. S6). A total of 30 units were identified as ‘tagged’ (IT^+^) with half-maximal inhibition occurring within 26 ms on average (median latency < 10 ms) (Fig. 3b, c, e). The mean duration of inhibition below the half-maximal level was 985 ms. Further consistent with selective identification, optotagged units were distributed at depths consistent with layer 5 (Fig. 3c).

**Fig. 3.**
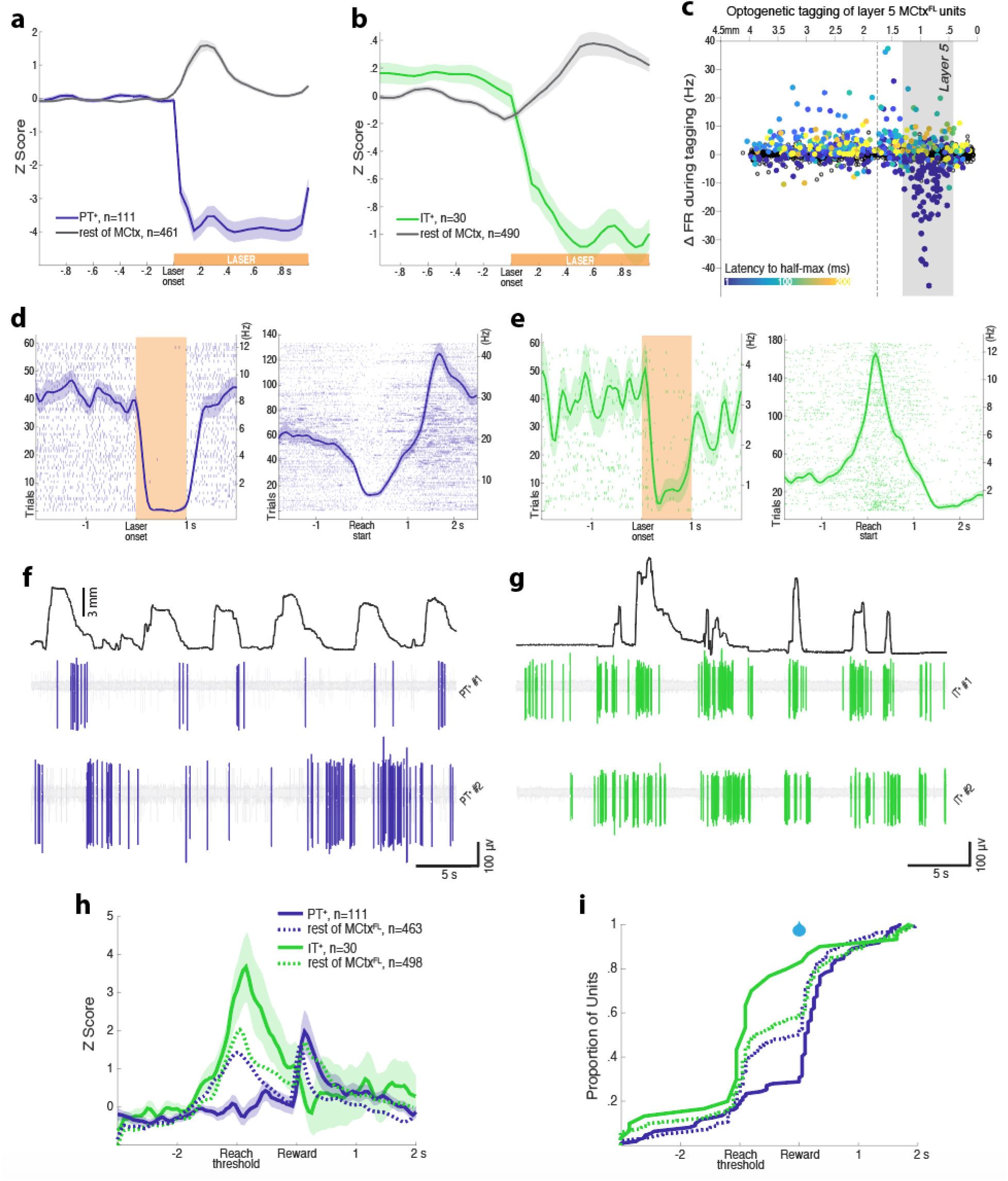
Prevalence of motor command-like activity in IT neurons. **a**, Normalized activity before and during optical silencing of pons-projecting PT^+^ neurons. **b**, same as **a** for striatum-projecting IT^+^ neurons. **c**, Mean firing rate change during optical tagging is plotted as a function of the inferred recording depth (x axis). Filled points indicate significant modulation. Latency to half-maximal firing rate change indicated with colorbar. **d**, *Left*, Activity of an example PT^+^ neuron to laser (60 trials, 594 nm) during optotagging (left) and aligned to movement start during task performance (right). More examples are shown in fig. S7. **e**, Same as d for an example IT^+^ neuron. **f-g**, Filtered, raw voltage traces showing spike activity of example PT^+^ and IT^+^ units with the amplitude of joystick movement plotted above. **h**, Normalized mean±SEM activity of PT^+^, IT^+^ neural populations aligned to joystick threshold crossing (ReachT) and reward delivery. The mean activity of the rest (untagged) of MCtx^FL^ is plotted in dotted curves for comparison. **i**, The cumulative distributions of the peak activity are plotted for PT^+^ and IT^+^ neural populations. Distributions of the rest (untagged) of MCtx^FL^ are plotted in dotted lines for comparison.

We next compared activity of these two populations of tagged, identified cell-types during task performance. Individual examples often revealed dramatic differences in the timing of activity across the two populations (Fig. 3d-g). As a population, the modulation of activity in PT^+^ population was more mixed as compared with the rest of the MCtx^FL^ populations during forelimb movement (Fig. 3h; group x time interaction; repeated measures ANOVA, F_1,40_ = 10.10, p = 4.66×10^-61^, main effect of group; ANOVA, F_1,572_ = 21.54, p = 4.30×10^-6^). In contrast, the IT^+^ population displayed a more consistent positive modulation of activity than the cortical population in general as well as than the PT^+^ population in specific (Fig. 3h; group x time interaction; repeated measures ANOVA, F_1,40_ = 7.78, p = 2.65×10^-42^, main effect of group, PT^+^ vs. IT^+^; ANOVA, F_1,139_ = 25.29, p = 1.49×10^-6^). The activity of the majority of PT^+^ neurons (69.4%) peaked after the reward delivery, while the majority of IT^+^ neurons (80%) were most active during movement initiation/execution before the reward delivery (Fig. 3i; 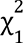 = 23.77, p = 1.08×10^-6^).

This difference appeared to be due at least in part to many pons-projecting PT^+^ neurons with suppressed activity around reach start (Figs. 3d, f, S7) similar to what has been described for the spinal-projecting subset of PT neurons previously (*40*). Thus, to compare similar activity patterns we sub-selected the PT^+,ext^ and IT^+,ext^ populations that displayed significant positive modulation of activity during movement (Fig. S8). We then examined the relative modulation of activity aligned to movement onset, movement offset, and threshold crossing/reward delivery. We found that positive modulation of activity, when apparent in both populations, appeared to be significantly greater in IT^+,ext^ than PT^+,ext^ regardless of temporal alignment (Fig. S8). The activity of PT^+,ext^ neurons often peaked after reward delivery in successful trials, whereas the activity of IT^+,ext^ neurons consistently peaked around reach start regardless of temporal alignment (Fig. S8; 7.09×10^-5^). 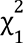 = 15.79, p = 7.09×10^-5^).

### Distinct, but complementary, movement kinematic encoding in Layer 5 IT and corticopontine PT neurons

We next asked whether identified Layer 5 corticopontine PT^+^ and corticostriatal IT^+^ neurons had similar or distinct correlates with movement kinematic parameters. We first examined the relative tuning of IT^+^ and PT^+^ neurons to movement amplitude and direction by comparing projections onto the AMP and DIR dimensions identified from simultaneously recorded population activity. A preponderance of IT^+^ neurons were strongly tuned to movement amplitude with modest or weak tuning to movement direction (Fig. 4a). In contrast, PT^+^ neurons showed the opposite propensity, with a majority more strongly tuned to movement direction than movement amplitude (Fig. 4a, right; KW test comparing relative tuning: p=3.0×10^-4^).

**Fig. 4.**
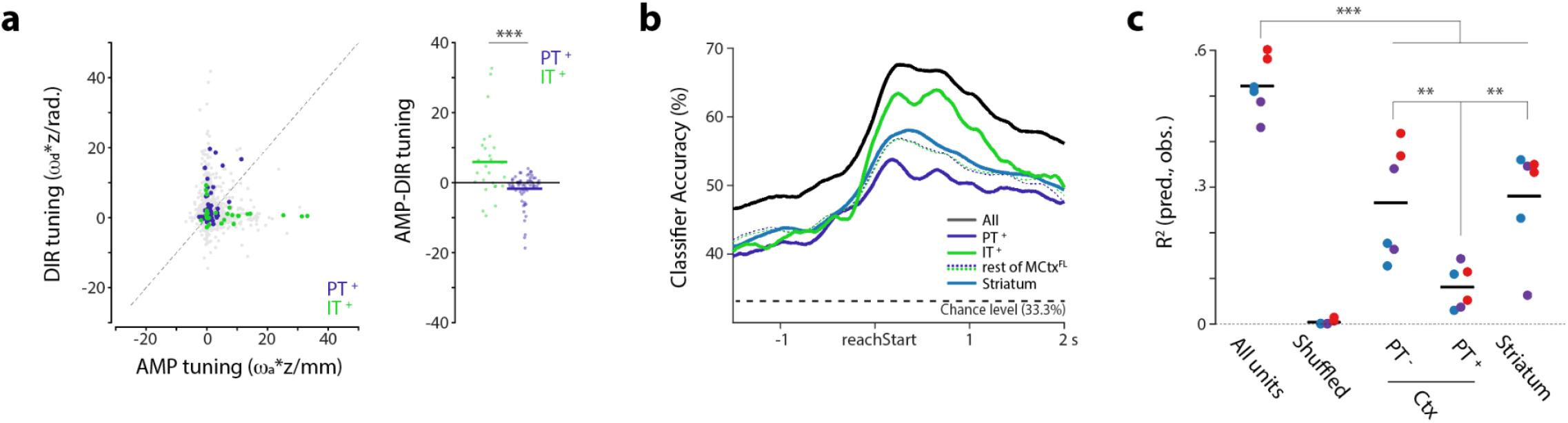
Projection cell classes are preferentially tuned to the amplitude or direction of forelimb movement kinematics. **a** *left,* Tuning of individual units to amplitude (x axis) and direction (y axis) plotted as slope of their weighted (AMP and DIR weights, ω_a_ and ω_d_ respectively) z-scored activity as a function of amplitude (units of mm) and direction (units of radians) quintiles. Full population of recorded units: grey; Optotagged, putative Sim1+ corticopontine PT (PT^+^, blue) and Tlx3+ corticostriatal IT (IT^+^, green) units highlighted. **a** *Right*, To compare preferential tuning across groups we compared the difference in tuning along AMP and DIR dimensions. Populations were significantly different (KW test, p<0.001). **b**, Cross validation performance of naive Bayes classifiers trained to predict movement amplitude tertile using all units (black) or optotagged populations (PT^+^, IT^+^) or striatum units as inferred from anatomical position. **c**, Contributions to committee decoder performance (see methods) for separate neural populations from recording sessions in Sim1-cre mice. Populations were identified by optotagging (PT^+^) or inferred from anatomical position are plotted; *\*\*\* p<0.001, ** p<0.01, * p<0.05*.

We performed a number of additional analyses to further confirm this difference in the encoding of movement kinematic parameters between IT^+^ and PT^+^ neurons. First, we confirmed that randomly chosen subsets of tagged IT^+^ neurons were indeed better at classifying observed movement amplitude when compared with PT^+^ neurons by training naive Bayes classifiers of movement amplitude (Fig. 4b). Given that our task was designed to vary substantially in movement amplitude driven by block wise changes in amplitude threshold, we reasoned that PT^+^ populations also carry relatively less information about overall movement kinematics in this task. We compared the decoding power for individual movement trajectories by assessing the relative contribution of PT^+^ neurons when compared to simultaneously recorded non-PT neurons (Fig. 4c). Again, we found that a relatively smaller amount of decoding information was present in PT^+^ neurons. Finally, we examined correlations between identified IT^+^ and PT^+^ individual neuronal activity and movement amplitude. Correlations were stronger in IT^+^ neurons than PT^+^ neurons even in the presence of variable movement direction and regardless of the sign of change in movement-related activity (Fig. S9; independent t-test, t_126_=3.65, p=3.82×10^-4^). Thus, across diverse analysis methods (preferential loading onto the AMP dimension of population activity, trialwise correlation of spike count, decoding, classifier performance) we observed consistent, significant evidence that corticostriatal Tlx3+ IT neurons encode information about the amplitude of movement when compared to corticopontine Sim1+ PT neurons.

### Two-photon calcium imaging of Layer 5 IT and corticopontine PT neurons

Identifying cell-types via optogenetic tagging has been an important technique that has clarified cell-type specific neuronal correlates; however, it is also subject to a number of limitations (*35, 50*). For example, some approaches such as antidromic stimulation are thought to have low false positive rates, but high false negative rates (*35, 54*). Whereas somatic stimulation (*50*) or somatic inhibition (used here) can have potentially higher false positive rates due to polysynaptic effects. Sustained inhibition (∼500 ms as used here) attempts to mitigate these false positives. Our best estimate of a putative false positive rate was ∼1% (see Methods) indicating that electrophysiological correlates in distinct cell types were likely mediated by true positives. Nonetheless, it is difficult to estimate such rates quantitatively without ground truth and thus we also sought to use a complementary method to assess the cell-type specific differences in the MCtx^FL^ neural correlates of forelimb movements. We used cell-type specific calcium imaging to more precisely target two major layer 5 neuron populations in MCtx^FL^. We used a virally-driven expression of GCaMP6f in Sim1-cre and Tlx3-cre mice (*35, 48*) as described for electrophysiological tagging experiments (Fig. 5a-b, fig S10; Sim1-cre: 8 mice, 19 imaging sessions, N=1576 ROIs. Tlx3-cre: 7 mice, 14 imaging sessions, N=1006 ROIs).

**Fig. 5.**
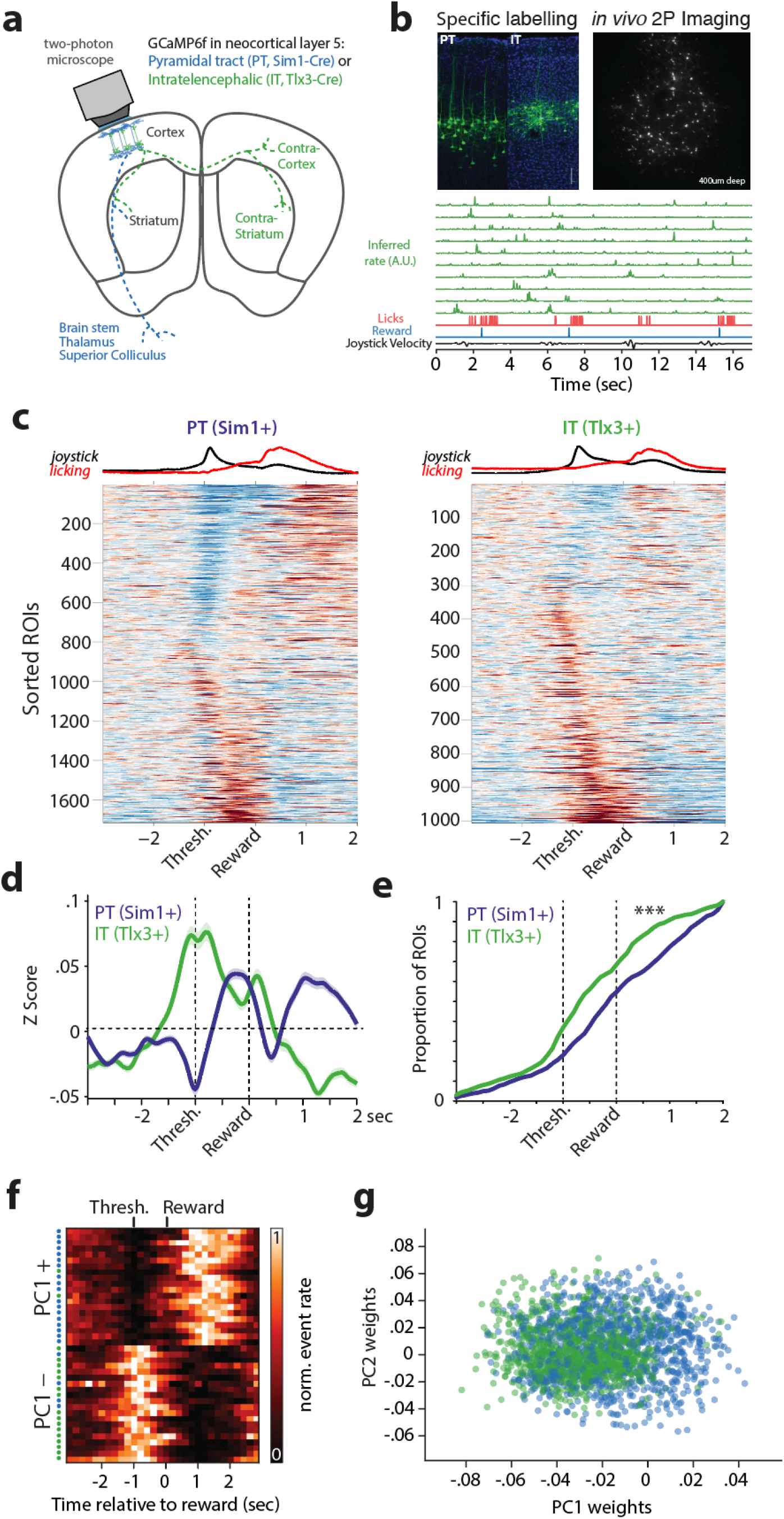
Calcium imaging shows cell-type specific differences in forelimb movement correlates. **a**, Two-photon calcium imaging schematic. **b**, *Top left*: Example histology from each mouse line. Scale bar 100 microns. *Top right*: Example imaging field of view. *Bottom rows*: Green traces are inferred spike rates of randomly selected IT neurons aligned to behavioral variables. **c**, Mean z-scored activity traces aligned to reward delivery for every imaged neuron (ROI) in the dataset. *Left,* PT (Sim1+); *Right,* IT (Tlx3+). ROIs are sorted by sign of movement-related modulation and time of peak modulation. Top row shows the average normalized joystick speed and lick rate. Scale: −0.5 (blue) to 0.5 (red) z. **d,** Mean activity for each cell-type. Shaded area is s.e.m. **e**, Cumulative proportion of maximal activity for each ROI (analogous to Fig. 3i). **f,** Normalized inferred spike rate for individual units with positive or negative PC1 loadings are plotted. Colored dots on the left reflect the cell type. For PCA, PT units were randomly subsampled to match the size of the IT population. Individual principal components, and additional example units, are provided in fig. S8. **g**, Histogram of unit weights on PC 1 for IT and PT neurons.

Imaging of pons-projecting PT neurons and layer 5a IT neurons showed prominent differences that were consistent with the electrophysiology data. IT neurons, similar to electrophysiologically recorded IT^+^ neurons, showed a bias towards larger peri-movement activation than PT neurons, while PT neurons had greater activation timed to reward consumption (Fig. 5c-e). The cell-type specific differences in imaging experiments were very similar to those observed with electrophysiological recordings (Fig. S12). To capture the variance in activity from the imaging experiments we examined population activity in a low dimensional state space spanned by the leading PCs. The first PC provided a dimension that distinguished activity of PT and IT populations. Cells with a negative loading on to the first PC (PC1−) were characterized by prominent activation around movement execution and were more likely to be IT neurons. In contrast, cells with a positive loading (PC1+) characterized by relatively suppressed activity during movement and more reward-timed modulation of activity were more likely to be PT neurons (Fig. 5f-g, fig. S10c-e, PT/IT difference on PC1: p < 5.35×10^-45^, independent t-test). Although these imaging analyses are consistent with prominent movement command-like activity in IT populations, we found that correlations with movement kinematics were detectable in both populations but were smaller and more variable in imaging data as compared to electrophysiological data (*55*)).

### Cell-type specific inactivation of IT and PT projection classes

We observed clear differences in the encoding of movement amplitude and direction in IT^+^ and PT^+^ neurons, respectively. If these differences reflect distinct, but complementary pathways by which descending motor commands influence movement it would predict dissociable effects on forelimb movements during inactivation of each cell type. In contrast, if IT neurons primarily exert their effects on movement through PT neurons, then we would expect similar or greater effects of PT^+^ inactivation as compared with IT inactivation (*35*). Thus, we next performed cell-type specific inactivation during movement execution with a potent optogenetic inhibitor (FLInChR (*53*)). To inactivate MCtx^FL^ populations during movement execution we used movement-triggered inactivation analogous to the pan-MCtx^FL^ activation (Fig. 1f). We used the same viral strategy with two mouse lines that restrict expression to (*2, 48*) layer 5 IT (Tlx-cre) and PT (Sim1-cre) neurons (Fig. S6).

Although there are clear limitations in attempts to directly compare perturbations across distinct mouse lines and experiments, it is important to confirm that inactivation produced a comparable change in the activity of target cell types. We triggered laser delivery at the earliest time point of reach initiation on a random subset (∼25%) of trials (schematized in Fig. 6a; also as in fig. 1f). We found that our perturbation strongly suppressed the PT^+^ neurons compared to their modulation during control movements (Fig. 6b, fig. S11a-b; ANOVA, F_1,220_=61.72, p=1.74×10^-13^). We next examined the magnitude of IT perturbation relative to its modulation during normal movements. We again observed a significant (but lesser in magnitude) inactivation (Fig. 6c, fig. S11c-d; ANOVA, F_1,58_=4.22, p<0.05). Thus, inactivation during movement is effective in both layer 5 corticostriatal IT and corticopontine PT neurons although there may be somewhat weaker inactivation of IT neurons.

**Fig. 6.**
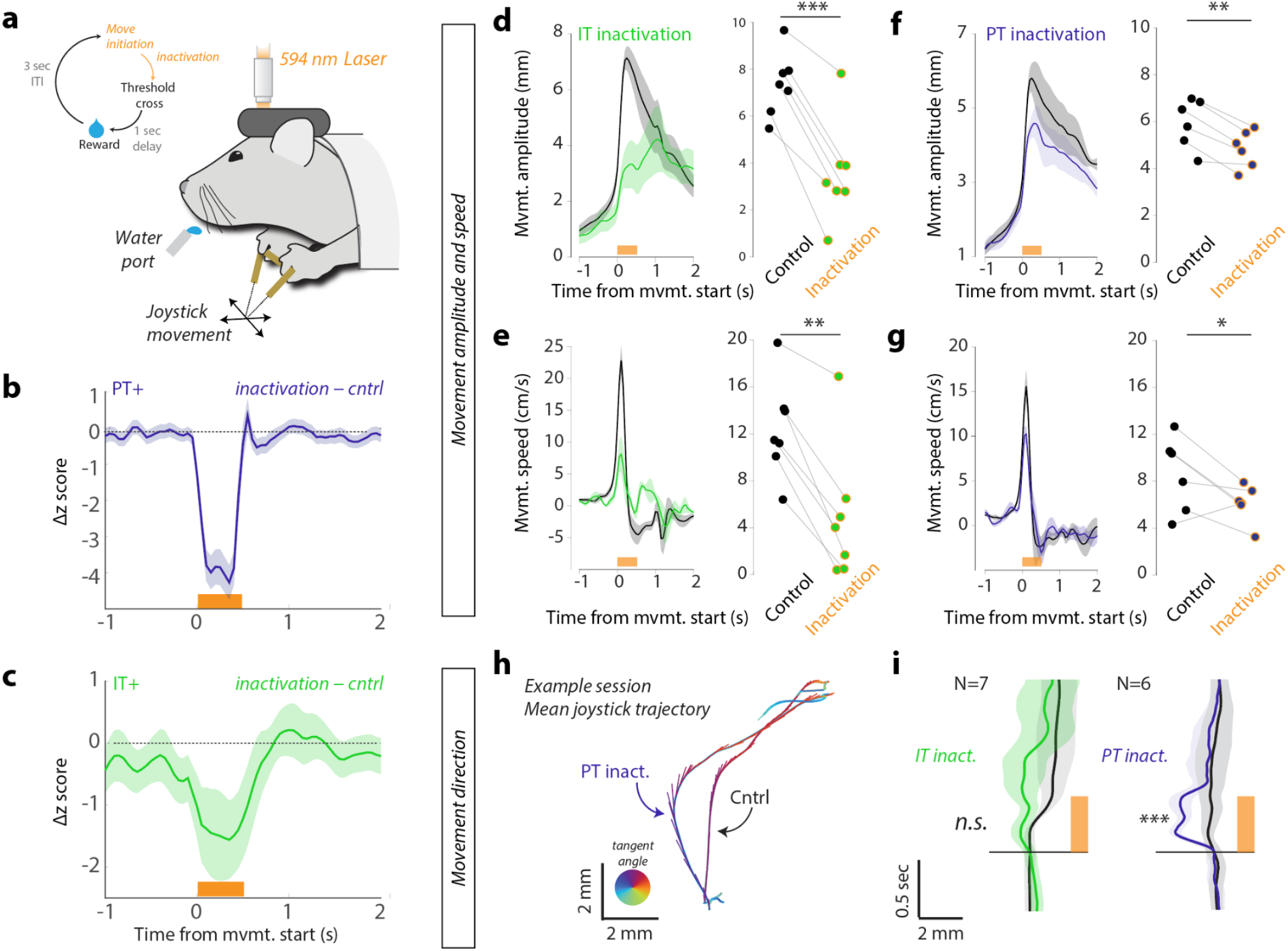
Differential effects of cell type specific optogenetic inactivation on forelimb movement kinematics. **a**, Schematic of closed-loop inactivation paradigm. **b**, Difference in movement-aligned activity between control trials and laser inactivation trials for identified corticopontine PT^+^ neurons. **c**, Difference in movement-aligned activity between control trials and laser inactivation trials for identified Layer 5 corticostriatal IT^+^ neurons. **d-i**, Behavioral effects of inactivation on movement amplitude and speed (**d-g**) and direction (**h-i**) were examined for inactivation of layer 5 IT neurons (**d-e**) and layer 5 corticopontine PT neurons (**f-g**). For each: *Left*, Mean±SEM reach amplitude/speed of unperturbed control trials (black) and perturbed inactivation (color) trials. *Right*, Mean reach amplitude/speed of unperturbed control (black dots) and inactivation trials (colored dots) for individual sessions. **h**, For an example session in Sim1-FLInChR mouse trajectories were reliably biased in direction on inactivation trials relative to control trials. Traces show mean trajectories with tangent vectors indicating speed (length) and direction of movement (angle). **i**, Population data showing x component of movement trajectory as a function of time for inactivation trials (IT, green; PT, blue) compared to control trials (black). Shaded area is s.e.m. *\*\*\* p<0.001, ** p<0.01, * p<0.05*.

To assess the relative contribution of these pathways to the execution of skilled forelimb movements we considered both effects on movement amplitude and speed (Fig. 6 d-g) and movement direction (Fig. 6h-i). Suppression of activity in MCtx^FL^ Layer 5 corticostriatal IT neurons led to a large reduction in movement amplitude and speed relative to control movements (Fig. 6d-e, paired t-test, amplitude: t_6_=8.13, p=1.85×10^-4^; speed: t_6_=5.26, p=0.002). Despite a larger inactivation, optogenetic suppression of corticopontine PT neurons led to a significant, but several-fold smaller effect on the amplitude (IT: −51±7%, PT: −19±2%; Mean±SEM percent reduction from the control trials) and speed (IT: −67±13, PT: −25±13) of forelimb movements in the skilled joystick movement task (Fig. 6f-g, paired t-test, amplitude: t_5_=6.55, p=0.001; speed: t_5_= 2.89, p=0.034).

Our previous analyses of neural correlates of movement indicated that corticopontine PT neurons may preferentially participate in the control of movement direction, at least relative to corticostriatal Layer 5a IT neurons. Thus, we next examined whether the trajectories of movements were altered during cell-type specific inactivation. Suppression of corticopontine PT neurons during movement elicited stereotyped changes in movement trajectory time-locked to inactivation that could readily be observed in single sessions (Fig. 6h) and a consistent directional bias in trajectories in all perturbation sessions (Fig. 6i; paired t-test, t_5_=7.26, p=7.73×10^-4^). In contrast, we found no clear change in movement direction when corticostriatal layer 5a IT neurons were suppressed during movement (Fig. 6i; paired t-test, t_6_=1.88, p=0.11). Thus, recordings from identified projection cell types revealed preferential encoding of movement amplitude in Layer 5 IT+ neurons and movement direction in corticopontine PT+ neurons. Inactivation of Layer 5 Tlx3+ IT neurons led to a preferential decrease in movement amplitude, and inactivation of corticopontine Sim1+ PT neurons preferentially altered movement direction.

### IT inactivation severely disrupts performance of skilled reach-to-grasp task

Analysis of neural correlates of movement in identified corticopontine PT^+^ and corticostriatal IT^+^ neurons and the effects of cell-type specific perturbation were very consistent in the joystick task. Specifically, corticostriatal IT neurons are preferentially tuned to movement amplitude and inactivation during movement reduces movement amplitude. Corticopontine PT neurons, in contrast, are preferentially tuned to movement direction and inactivation altered movement direction with smaller effects on movement amplitude. It has also been proposed that the role of motor cortical projection cell types could critically depend upon the task context or problem. Thus, we next asked whether these effects, in particular a critical role for the general IT class of projection neurons, are specific (limited to) the highly variable joystick movements. For example, it has often been proposed that dexterity demands could be intimately tied to the function of corticopontine PT pathways (*4*) and yet inactivation of dorsal striatum also profoundly impairs movement amplitude during reach-to-grasp tasks (*21*) as it does for joystick movements (*22*) perhaps consistent with a role for corticostriatal IT neurons. Moreover, inactivation of the basal pons produces little effect on movement speed and amplitude (*56*).

Thus, we next examined whether IT neurons are critical for the execution of forelimb movements in a head-fixed reach-to-grasp task for mice (*44, 57*) (Fig. 7a, Supplementary Video 2-3). To assess whether there was any potential role for IT neurons we adopted a penetrant strategy and labeled IT neurons just with contralateral retrograde AAV injection (more similar to labeling approaches in previous studies (*36*)), and the corticopontine PT neurons were labeled with retrograde AAV injection into the basal pons. We first confirmed that both strategies were sufficient to produce robust inhibition of layer 5 IT and PT neurons (Fig. 7b-c). Next, we delivered laser (2 sec) at the presentation of the food pellet in randomly selected trials, and examined how the cell-type specific inactivation affected animals’ movement and overall performance. In some trials, the inactivation was triggered on initial movement of the forelimb off its resting position and towards the food pellet (Supplementary Video 2). Similar to our results in the variable amplitude and direction joystick task, we found that silencing IT neurons led to profound disruptions in reach-to-grasp performance by dramatically attenuating reach movement amplitude and blocking progression to next movement components (Fig. 7d, Supplementary Video 2; paired t-test on success rates, t_7_=18.91, p=2.88×10^-7^). These effects were similar to previously described impairment of forelimb movements in this task with pan-cortical inactivation (*44*). In contrast, silencing corticopontine PT neurons did not impair movement amplitude and performance quality was largely intact (Fig. 7e, Supplementary Video 3; paired t-test on success rates, t_7_=0.57, p=0.58). Together, these data provide strong evidence that IT projection neurons play a key role in reach-to-grasp behavior that is not fully explained by an IT through corticopontine PT pathway.

**Fig. 7.**
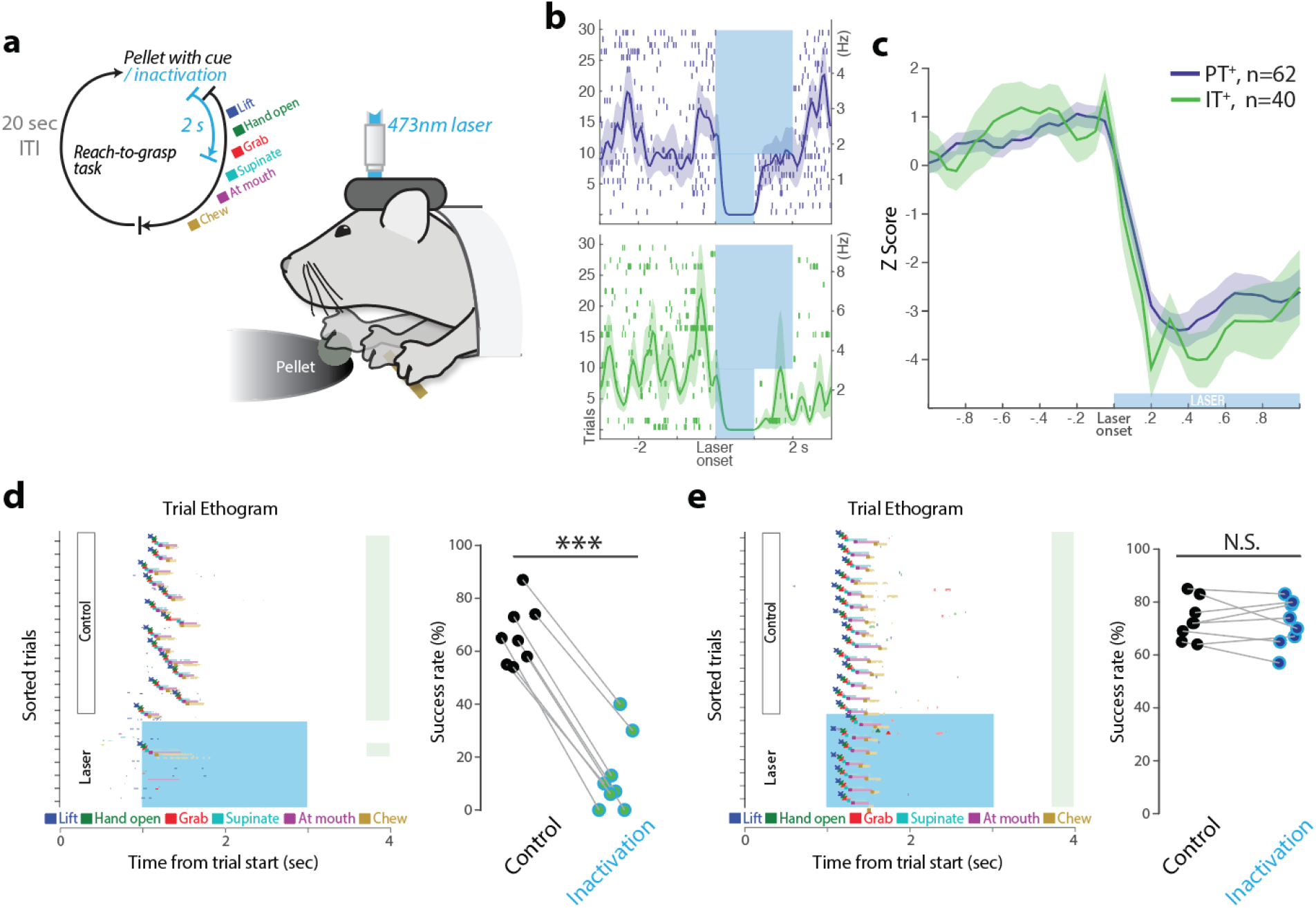
MCtx^FL^ IT neurons are necessary for execution of forelimb movements in a reach-to-grasp task. **a**, Schematic of inactivation paradigm in reach-to-grasp task (*57*). Laser was triggered in randomly selected trials (∼27%) at the cue onset. **b***, upper*, representative PT neuronal response to optogenetic inactivation with GtACR2. Lasers were delivered for 1 s or 2 s in interleaved trials. *lower*, An IT neuronal response to inactivation. **c**, Identified PT^+^ units (62 from 4 mice) and IT^+^ units (40 from 4 mice) displayed comparable responses to optical silencing. **d**, *Left*, Inactivation of IT neurons in the contralateral (left) hemisphere of the reaching arm (right) blocked initiation and successful execution of reach-to-grasp presented as an ethogram of a representative session with each behavioral component automatically labelled by JABBA (*74*). *Right*, Fraction of successful performance in control versus IT inactivation trials for all sessions (n=4 mice, 8 sessions). **e**, *Left*, Inactivation of PT neurons had no effect on task performance in a representative session. *Right*, Fraction of successful performance in control versus PT inactivation trials for all sessions (n=4 mice, 8 sessions).

### Inactivation of PT and IT neurons oppositely affect striatal activity

Inactivation of Layer 5a IT neurons produced substantial reductions in movement amplitude without substantial alterations in movement direction consistent with preferential encoding of movement amplitude in this population. This effect was also quite distinct from PT inactivation which produced smaller changes in movement amplitude and a clear change in movement direction - consistent with a preferential encoding of movement direction in the Layer 5b corticopontine PT population. This is more consistent with complementary, but separate descending pathway from corticostriatal Layer 5a IT neurons that determines movement amplitude, as compared with the proposal that IT output may exert its effects on movement execution through an intracortical IT→PT pathway (*4, 35*). Since the only extra-cortical output of IT neurons is striatum (and in the case of MCtx^FL^ primarily dorsal striatum), this suggests that the effects of inactivating corticostriatal Layer 5a IT and corticopontine Layer 5b PT would also differ in their consequences for activity in dSTR.

Previous work with retrograde labeling indicates that a given region of dSTR receives at least partially convergent input from IT and PT neurons within a given cortical column (Fig. 8a) (*3, 29, 52*). Thus, we finally sought to assess whether inactivation of IT^+^ and PT^+^ had differential effects on striatal activity or a largely shared effect as expected if mediated primarily by an intracortical IT→PT pathway. Optogenetic suppression of IT neuron activity during movement results in a corresponding decrease in forelimb movement-related striatal activity consistent with IT providing a source of direct excitatory input (Fig. 8b). However, during optogenetic silencing of pons-projecting PT neurons we found that striatal units on average *increased* activity during PT inactivation and that this differed significantly from IT inactivation (Fig. 8b; striatal modulation by PT vs IT inactivation; ANOVA, F_1,1146_=35.49, p=3.41×10^-9^). Striatum is composed of a number of cell types including both inhibitory projection neurons (medium spiny neurons; MSNs) and local interneurons. Although it is not possible in these datasets to distinguish cell types based upon molecular identity, as with other brain regions these two broad classes are roughly distinguished by their baseline firing rates. PT inactivation resulted in small increases in the activity in the subset of striatal units with low (<10 Hz) baseline firing rates - including MSNs (Fig. 8c; ANOVA, F_1,972_=4.41, p=0.03). In contrast, we again observed a differential consequence of IT inactivation reflected in robust decreases in the activity even for the subset of neurons with relatively low firing rates (Fig. 8d; ANOVA, F_1,780_=8.29, p=0.004).

**Fig. 8.**
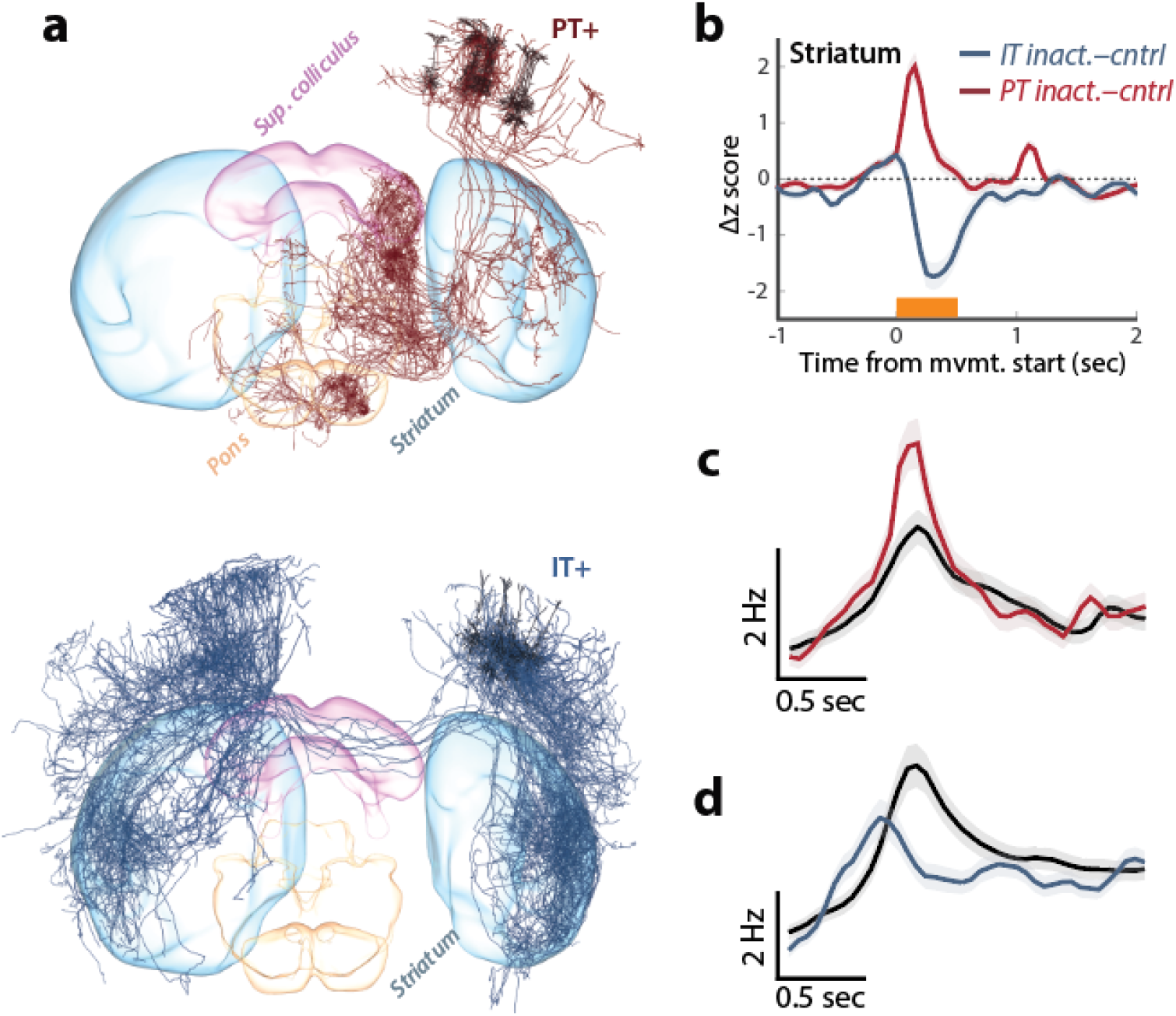
Inactivation of PT and IT neurons oppositely affect striatal activity. **a**, 3D visualization of complete single neuron reconstructions (*3*) for 10 representative single cell reconstructions of layer 5 PT (top) and IT (bottom) anatomical classes from the mouselight.janelia.org database show partially overlapping projections to the recorded region of dorsal striatum (dStr). **b**, For all units from dStr we computed the difference between movement aligned, z scored time histogram for control trials and perturbation trials in which either Sim1+ corticpontine PT neurons (red) or Layer 5 Tlx3+ IT neurons (blue) were inactivated during movement. **c-d**, Populations of units with low average firing rates (also broad spike widths on average) were used to assess whether modulation of dStr activity was consistent with changes in medium spiny projection neuron activity during PT (c) or IT (d) inactivation trials. Control trials (black) reveal clear movement-aligned modulation of activity in these populations and opposing changes during inactivation.

## Discussion

The central control of movement is characterized by the ability to execute movements adapted to achieve diverse goals with a common effector. For example, animals can use their forelimbs over a continuously varying range of speed and amplitude (*26*), utilize one or both forelimbs in a coordinated fashion (*58*), targeted to variable manipulanda (*59*), or reach out to eccentric targets in a range of directions (*44*). The circuit mechanisms that allow the same putative motor cortical circuits to control these movements and learn to adapt specific components during the development of motor skills has been difficult to understand. However, a division of computational labor across distinct anatomical loops spanning the motor cortex through the brainstem is thought to be critical (*10*). In particular, two largely (although not completely) distinct cortical-subcortical circuits - the basal ganglia and cerebellum - have long been thought to play complementary, but distinct roles in the control of forelimb movements (*10, 11*). The extent to which the differential functions of cortico-cerbellar-thalamic and cortico-basal ganglia-thalamic loops are due to differences in the cortical projection neuron classes they involve is unclear.

Many studies of cortical encoding of movement have focused on cued movements of individual limbs with repeatable and stereotyped trajectories, often in multiple directions, but with relatively little variation in movement speed/amplitude. Here we used a paradigm in which mice vary movement amplitude over roughly an order of magnitude and explore a range of movement directions (albeit with relatively little lateral movement). Using this dataset we discovered two orthogonal dimensions of population activity that explained much of the variance in movement amplitude and direction. We found that movement correlates in Layer 5a corticostriatal IT and Layer 5b corticopontine PT projection neuron classes were not homogeneously distributed, but rather preferentially participated in the amplitude and direction dimensions of population activity, respectively. These data are consistent with prior observations of preferential tuning of PT neurons to movement direction in mice (*35*) and in primates (*60*), but also highlight how consideration of another key aspect of movement kinematics - amplitude and speed - can reveal further complexity in cell-type specific components of motor cortical activity.

Here, we focused on two broad classes of molecularly and anatomically distinct cortical Layer 5 projection neurons - corticostriatal IT and corticopontine PT (*1*). PT neurons are a diverse class composed of multiple anatomical and molecular subtypes (*2, 3, 9*). Corticopontine PT neurons are the presumptive source of descending motor command information used by the cerebellum to compute forward models (*20*). We explored forelimb tasks with more diverse movement kinematics and found that deep layer 5 neurons and in particular a subset of corticopontine PT neurons were preferentially tuned to movement direction, consistent with preferential direction tuning in pons-projecting primate PT neurons (*41*). At the same time it remains possible that with more diverse classes of forelimb movement (e.g. grabbing multiple objects) and finer cell-type information, more complex cell-type specific tuning will be discovered. Our data suggest that as movement variability increases more distributed, cell-type specific activity is revealed; reflecting the fact that task design itself is a key determinant of the observed dimensionality of motor cortical activity (*61*).

Our data are potentially surprising from a perspective in which pyramidal tract pathways are a primary determinant of descending cortical influence on motor commands through projections to spinal cord and other subcortical areas (*4, 62*). At the same time, our data are broadly consistent with movement execution-related activity observed in multiple cortical cell types in rodents (*63*) and primates (*8, 41*). Recent work in primates with antidromic identification of projection types and a forelimb movement around a single (elbow) joint, found lesser tuning of corticostriatal neurons to movement kinematics as compared with pyramidal tract neurons (*41*). This may reflect a difference between mice and primates, but also may be a consequence of different movement dimensionality, analytic techniques, or different sampling biases. Pasquerau *et al.* (*41*) used antidromic activation in the posterior putamen to identify corticostriatal neurons that could potentially sample from more numerous superficial IT neurons (lamina in which we also tended to find weaker decoding and correlates with movement amplitude; Fig. 2f). Similarly, it is unclear whether our labelling strategy for corticopontine neurons was biased towards a distinct subset of PT neurons relative to previously studied populations. Lastly, here we developed distinct methods to find orthogonal coding dimensions and decode movement kinematics in much more variable movements that will be intriguing to use in future work on primate datasets and further understand these differences.

It has long been proposed that multiple, parallel ‘modules’ in the brain may be critical for flexible control of movement (*18*). While these parallel modules (potentially, forward/inverse models) are thought to be instantiated in cerebellum (*20, 64*), the division of labor across corticostriatal IT and corticopontine PT cell types into distinct roles controlling movement amplitude and direction, respectively, is consistent with the dissociable effects of basal ganglia and cerebellar perturbations. The effects of cerebellar perturbation have tended to be primarily deficits in either the targeting of movements in a specific direction or adaptation of movement direction to an environment change (e.g. visuomotor rotation) (*20*). In contrast, perturbations of basal ganglia function often lead to aberrant control of movement amplitude and speed (*22, 24, 26, 30, 65*). Basal ganglia pathways have been proposed to control movement amplitude/speed either by adaptively adjusting the gain of motor commands based upon reward feedback (referred to as movement vigor (*26, 27, 33*)) or by determining a reference signal for a continuous feedback controller (*31*) or by producing motor commands *per se* (*32*). In the context of the current experiments these models all make similar predictions and thus cannot be distinguished in detail, but are broadly consistent with a pathway involving corticostriatal IT and dSTR neurons being a critical “module” involved in descending forebrain control of movement amplitude. dStr also receives input via collaterals of corticopontine PT neurons. The extent to which these pathways are kept separate or potentially re-integrated in subcortical target areas will be a key question in future work.

Circuit mapping experiments have revealed an asymmetric IT→PT excitatory connectivity (*36*). On the one hand, the absence of strong PT→IT connectivity could help to explain how IT activity is not strongly tuned to movement direction. On the other hand, it is less clear how prominent movement amplitude correlated activity in IT neurons is relatively weak or not present in corticopontine PT neuron activity. IT neurons project onto other (e.g. corticospinal (*36*)) PT populations not studied here and one possibility is that non-corticopontine PT neurons are the primary recipients of preferential IT input, indeed previous work focused more on retrogradely-labelled corticospinal PT populations and cross-hemisphere IT neurons - both of which are subsets with different biases in labelling compared to the methods used here (*36*). Although pons-projecting PT neurons are a large subset of PT neurons, our approach was likely not penetrant for all subclasses of PT neurons (*2, 3, 48, 66, 67*). The molecular marker Sim1+ may bias against corticospinal and corticothalamic PT neuron classes (*2*). Future work further dissecting molecular subtypes of PT neurons (*54*) in connection with detailed information about the cortical microcircuit in which those neurons are embedded (*36, 68, 69*) will be critical to understand how descending output is distributed across projection classes. Another possibility is that the extensive PT dendrites (relative to IT) which can powerfully modulate the integration of multiple sources of afferent input (*70, 71*) could be critical to transform movement amplitude-biased activity in IT neurons to movement direction-biased activity in corticopontine PT neurons - perhaps via gating by another input to corticopontine neurons related to muscles group identity (flexor vs extensor). Such a model also has the merit of potentially providing flexibility - if experience led to a learned reduction in the putative gating input for direction, corticopontine PT neurons could then also be correlated with movement amplitude. Clearly, this remains a speculative hypothesis that will require additional studies in animals learning multiple motor tasks to resolve, but existing work suggests the possibility of dramatic remapping of the relationship between PT activity and movement can occur (*62*).

The discovery that direct motoneuron innervation by PT corticospinal neurons is unique to primates has provided an anatomical justification for accounts in which PT projections are particularly central to the remarkable motor skills of primates relative to other mammals (*4, 42*). However, increasing diversity of IT neuron populations is also a primate innovation (*9*) and thus correlated with increasing flexibility of motor skills. We note that basal ganglia output plays a role in controlling movement speed and amplitude in human and non-human primates (*26, 28, 72*) as it does in other mammals, e.g. mice (*27*). Although less well studied in the context of the control of movement execution, these considerations suggest that it will also be critical to study non-PT motor cortical projection cell types which may provide new insights into motor cortical function in diverse species. Here we describe approaches that allow robust single trial decoding of movements and an approach using targeted dimensionality reduction to identify independent components of population activity that encode separable parameters of movement kinematics. This approach may prove useful for future models of descending motor commands that are distributed across cortical projection classes and subcortical target areas.

## MATERIALS AND METHODS

Male and female mice, typically aged 8-16 weeks at time of surgery, were used in this study. All procedures were approved by the Janelia Research Campus Institutional Animal Care and Use Committee (IACUC) and were consistent with the standards of the Association for Assessment and Accreditation of Laboratory Animal Care. Mice were water restricted (1-1.2ml water/day), and their weight and signs of health were monitored daily as in(*22*).) Surgical methods closely followed those previously described(*22, 73*) except where indicated below.

### Behavior

#### Joystick task

The variable amplitude operant task was run as described previously (*22*) using a microcontroller based system (details can be obtained from http://dudmanlab.org/html/resources.html). After surgery (see below), mice were given 5 days of recovery prior to beginning water restriction (1ml water/day). Following 3-5 days of initial water restriction, they underwent 10-20 days training, which simply involved exposure to the task and self-learning. Mice were head-fixed in a custom made head restraint box using the RIVETS head-fixation apparatus (*73*). The mouse’s front paws rested on a metal bar attached to a spring-loaded joystick, which had unconstrained 2D maneuverability in the horizontal plane. Mice were trained to maneuver the joystick to certain thresholds varying across three different blocks (e.g. 4.2-5.7-4.2 mm) to obtain a sweetened water reward delivered 1 s after each threshold crossing. Rewards were followed by a 3 s inter-trial interval (ITI) in which no movements would be rewarded. There were up to 150 trials (50 trials per block) in electrophysiology and 120 trials per session in imaging (some sessions were incomplete), with one water reward being available per trial. All behavioral events (forelimb movements, licks) were recorded on separate channels at 25 kHz (USB-6366; National Instruments, Austin, Texas) then downsampled offline at 1 kHz. Forelimb movements were assessed offline to detect individual reaches based on the speed joystick movement. Time points of reach start and stop were defined as well as other kinematic properties such as duration, maximum amplitude and speed for each reach.

#### Reach-to-grasp task

Methods are as described previously (*44*). Briefly, mice were habituated to head fixation (*73*) in a light tight, ventilated, soundproof 28-inch cubic behavioral box. Mice were initially trained for approximately 30 min per day, until they started licking pellets (10 or 20 mg; Test Diet; St Louis, MO) placed directly below their mouth. Food pellets arrived ∼ 200 ms after the start of an auditory tone (5 kHz) by rotating the turntable with a servomotor driven by custom-programmed Arduino software. Mice were initially (1–5 training sessions) trained to retrieve a food pellet by licking and eating the pellet, often using their hand to guide the pellet into their mouth. After cued licking was learned, the turntable was moved progressively further away (over 3–10 sessions) to encourage mice to reach for the pellet after the cue. Mice almost always started with hands on perch and trials where animals lifted the hand before the cue were discarded. Mice were trained each day for approximately 60 min until they routinely responded to the auditory cue (within 1 s) and grabbed the pellet.

Two high-speed, high-resolution monochrome cameras (Point Grey Flea3; 1.3 MP Mono USB3 Vision VITA 1300; Point Grey Research Inc.; Richmond, BC, Canada) with 6–15 mm (f/1.4) lenses (C-Mount; Tokina, Japan) were placed perpendicularly in front and to the right of the animal. A custom-made near-infrared LED light source was mounted on each camera. Cameras were synced to each other and captured at 500 frames/s at a resolution of 352 × 260 pixels. Video was recorded using custom-made software developed by the Janelia Research Campus Scientific Computing Department and IO Rodeo (Pasadena, CA). This software controlled and synchronized all facets of the experiment, including auditory cue, turntable rotation, optogenetic lasers, and high-speed cameras. Fiji video editing software was used to label laser onset, termination, and timestamp in the videos. Annotation of behavior was accomplished using JAABA (*74*) as described previously (*44*). See supplemental videos for examples of individual trials and effects of inactivation.

### Extracellular electrophysiological identification and recording of PT and IT neurons in awake head-fixed mice

For cell-type specific in vivo recordings from motor cortex and striatum in mice performing the variable amplitude operant task, rAAV2-retro-CAG-Flex-FLInChR-mVenus (3.0E+12 GC/ml) was injected to the pons bilaterally (relative to lambda: 0.4 mm anterior, 0.4 mm lateral, 5.5, 5.75, 6 mm deep, 70 nL/depth) in Sim1-cre (KJ18Gsat RRID:MMRRC_037650-UCD) mice, selectively labeling a pyramidal type (PT) layer 5 population (*48, 52, 54*). The same viral vector was injected to the dorsal striatum (relative to bregma: 0.5 mm anterior, 1.6 mm lateral, 2, 2.7, 3.5 mm deep) and cortex (site 1 : 0.9 anterior, 1.5 lateral, site 2: 0.1 anterior, 1.9 lateral, site 3: 0.1 anterior, 1.1 lateral, each site at 0.3 and 0.6 mm deep, 80nl/depth) in Tlx3-cre (PL56Gsat RRID:MMRRC_041158-UCD (*48*)) mice, selectively labeling a layer 5 IT population. Prior to recordings, a craniotomy was made over the recording sites (relative to bregra: 0.5 mm anterior, ±1.7mm lateral) at least 12 hours prior to recording under isoflurane anaesthesia. Exposed brain tissue was kept moist with phosphate-buffered saline at all times, and craniotomy sites were covered with Kwik-Sil elastomer (WPI) outside of the recording session.

For neural population recording during joystick behavior using the Neuropixels probe (*49*), awake mice fully recovered from craniotomy were head-fixed in a RIVETS chamber (*73*). A Neuropixels probe (option 3 phase A) with 374 recording sites was briefly (∼2 minutes) dipped into the diI cell-labeling solution (ThermoFisher) to visualize probe tracks, then lowered through the craniotomy manually. After a slow, smooth descent (0.2 mm/min), the probe sat still at the target depth for at least 5 min before initiation of recording to allow the electrodes to settle. An Ag wire was soldered onto the reference pad of the probe and shorted to ground. This reference wire was connected to an Ag/AgCl wire was positioned on the skull. The craniotomy and the Ag/AgCl wire were covered with a saline bath. Voltage signals are filtered (high-pass above 300 Hz), amplified (200x gain), multiplexed and digitized (30 kHz) on the base, allowing the direct transmission of noise-free digital data from the probe, and were recorded using an open-source software SpikeGLX (https://github.com/billkarsh/SpikeGLX). Recorded data were pre-processed using an open-source software JRCLUST (https://github.com/JaneliaSciComp/JRCLUST) to identify single-units in the primary motor cortex (M1) and STR. To assay FLInChR expression and responses, a fiber (200 mm core, 0.39 NA, Thorlabs) coupled to a 574 nm laser source (Omicron) was placed to deliver light onto the craniotomy. Single laser pulses of 1 s duration with power measured at the tip of the fiber of 4-8 mW were delivered 60 times with 8 s intervals. Mice were at rest after task completion during tagging.

For cell-type specific recordings from motor cortex in mice performing the reach-to-grasp task, rAAV2-retro-hSyn-GtACR2-KV-eGFP (8.5E+13 GC/ml) was injected to the pons bilaterally (relative to lambda: 0.4 mm anterior, 0.4 mm lateral, 5.5, 5.75, 6 mm deep, 30 nL/depth) in Sim1-cre (KJ18Gsat) mice, selectively labeling a pyramidal type (PT) layer 5 population. The same viral vector was injected to the dorsal striatum (relative to bregma: 0.5 mm anterior, 1.7 mm lateral, 2.8, 2.6, 2.4 mm deep) and cortex (0.5 mm anterior, 1.7 mm lateral, 1.0, 0.5 mm deep) in the right hemisphere of Slc17a7-cre mice to selectively label a layer 5 IT population in the left hemisphere. All recordings and optical silencing were conducted in the left hemisphere contralateral to the reaching hand (right). In the subset of mice used for reach-to-grasp task neural recordings to confirm inactivation were targeted to Layer 5 neurons using silicon probe arrays as described previously (*57*). A unit with a significant reduction in the spike count during the laser (paired t-test, α=0.01 and/or at least 60% reduction or 0.3 z-score sustained throughout laser pulse) relative to the baseline period) was considered to be optogenetically tagged. There was no difference between stringent and lenient tagging estimates for IT neurons, but there was a difference in PT neurons. To estimate the false positive rate we used the anatomical distribution of tagged PT units. This analysis yielded a false positive estimate of 0.3% (stringent) and 1.5% (lenient); see Fig. S12. We used stringent criteria for analyses that depended upon single cell properties (Fig. 4a) and lenient criteria when the maximal possible contribution of PT was considered (Fig. 4b-c) or population averages over the entire sample were compared (Fig. 3). Moreover, there is a good correspondence between population mean profiles obtained via optogenetic tagging and cell-type specific imaging (Fig. S12). These estimated false positive rate estimates suggest (1) conclusions are almost exclusively drawn from true positives and (2) we are likely still in a false negative dominated regime as expected for optogenetic tagging.

### Cell-type specific closed-loop perturbation of M1 neuronal activity

To examine the cell-type specific role of the deep layer 5 PT neurons in MCtx, we injected rAAV2-retro-CAG-Flex-FLInChR-mVenus(*52, 53*) into the pons (relative to lambda: 0.4 mm anterior, 0.4 mm lateral, 5.5, 5.75, 6 mm deep, 70 nL/depth) in three Sim1-cre (KJ18Gsat (*48*)) mice. Viruses obtained from Janelia Viral Tools (https://www.janelia.org/support-team/viral-tools). To examine the role of the IT neurons in MCtx, we bilaterally injected the same virus into the dorsal striatum (relative to bregma: 0.5 mm anterior, 1.6 mm lateral, 2, 2.7, 3.5 mm deep, 150 nL/depth) and cortex (site 1 : 0.9 anterior, 1.5 lateral, site 2: 0.1 anterior, 1.9 lateral, site 3: 0.1 anterior, 1.1 lateral, each site at 300+600 microns deep, 80nl/depth) in five Tlx3-cre (PL56Gsat (*48*)), respectively. In closed-loop experiments, a 500 ms single pulse of 574 nm laser was delivered bilaterally in randomly selected 30 % of the trials immediately when mice moved the joystick by 1.5mm from the zero point taken at the end of each ITI.

To examine the general role of MCtx in control of forelimb movement regardless of the projection neuronal cell-type, we implanted optical fibers (200 mm core, 0.39 NA, Thorlabs) bilaterally to place fiber tips right onto the pia of the brain in VGAT-ChR2-eYFP (*75*) (Fig. 1f-g) or Rbp4-cre RRID:MMRRC_037128-UCD (*48*)::Ai32 RRID:IMSR_JAX:024109(*76*) (Fig. S1) mice. In closed-loop experiments, a 500 ms single pulse of 473 nm laser was delivered in randomly selected trials triggered by a slight joystick movement caused by mice. In open-loop experiments, a 3 s single pulse of 473 nm laser was delivered in randomly selected 30 % of trials at a given time point (2 s after previous reward delivery during inter-trial interval in select trials) regardless of animals’ behavior.

### Cell-type specific two-photon calcium imaging

Viruses were AAV 2/1-Flex-GCaMP6f, diluted to 2*10^12^gc/ml (*77*) RRID:Addgene_58514 and obtained from Janelia Viral Tools (https://www.janelia.org/support-team/viral-tools). 5 injections performed in a cross-shape, centered on 1.6 lateral, 0.6 rostral. 20nL was ejected at 600um depth. This center was chosen based upon previous microstimulation work (*78, 79*). Imaging was restricted to one month after injection to minimize overexpression.

3mm-wide circular imaging windows were made over the left cortical hemisphere in all animals, following the method of Goldey *et al* (*80*). Window implants were centered on the virus injection center, and fixed in place using cyanoacrylate glue and dental acrylic. Windows (custom ordered from Potomac photonics) were made by placing three windows together, with the top window being 3.5mm, the bottom two being 3mm, such that the top window rested on a thinned skull area. This triple window arrangement was used to increase downward pressure on the brain and stabilize the brain motion.

Imaging was performed with a custom built two photon laser scanning microscope running scanImage software (latest versions, from 2013-2016; https://vidriotechnologies.com). GCaMP6f was excited with a ti:sapphire laser, tuned to 920nm. Imaging was typically performed at 33Hz via bidirectional scanning with a resonant galvo. Power at sample did not exceed 150mW. In poorer quality windows, frame rate was halved to allow an increase in peak pulse power. This was done to minimize photodamage from thermal effects. Depth of recording ranged from 350um-450um, depending upon imaging clarity, corresponding to the proximal dendritic region of the apical dendrite.

All imaging data analysis was performed in Python using custom-written scripts unless otherwise stated. Imaging data was motion corrected in two stages. Firstly, an image average was taken for a session across all frames. Secondly, each frame was then motion registered to that image, based upon a Fourier-based cross-correlation approach to detect the optimal corrective displacement. The average was then re-taken, and the process repeated 3 times. The result of this image registration process was examined by eye for each session to check for errors.

Region of interest (ROI) extraction was done manually in imageJ software. ROIs with high baseline fluorescence, a putative marker for unhealthy cells, were not used. Fluorescence traces were deconvolved to inferred rates using published code (*81*). We note that this is not an attempt to claim specific firing rates of neurons, but rather to reduce the distorting effect of the calcium sensors’ slow kinetics on the inferred activity. We did not attempt to calibrate these inferred spike rates with real rates.

### Histology

#### Fluorescence light sheet microscopy of cleared mouse brain

At completion of all electrophysiological experiments, mice were perfused with 40 ml of cold PBS (pH 7.4) containing 20 U/ml heparin at ∼10ml/min, and fixed with cold 4% PFA. Extracted brains were further fixed for 24hrs in 4% PFA. Fixed brains were delipidated using the CUBIC-L cocktail 10 w%/10w% N-butyldiethanolamine/Triton X-100 for a week. Delipidated brains underwent nuclear counterstaining with TO-PRO-3 (ThermoFisher) for a day. We then transparentized the delipidated brains in the refractive index (RI) matching cocktail CUBIC-R composed of 45 w%/30 w% antipyrine/nicotinamide for two days (*82*). Finally, cleared brains were imaged using fluorescence light sheet microscopy (Zeiss Lightsheet Z.1) to visualize expression of FLInChR (509 nm), probe tracks (570 nm), and nuclear counterstaining (661 nm).

The imaged 3D brain volumes (v3D) were aligned to a standardized brain coordinate system (Allen Anatomical Template, AAT) using a semi-manual landmark-based method (big warp) (*83*). The v3Ds were additionally aligned to the template MRI image volume (MRI3D) acquired using fixed brains in the skull to further correct for any distortion due to extraction of the brain from the skull (*84*). Each probe track was manually marked on v3D fused with AAT, and the 3D coordinates of all electrode sites were finally determined on MRI3D using the mapping between AAT and MRI3D combined with the geometry of the Neuropixels probe. Using the 3D coordinates, each electrode site was labeled as a brain region according to AAT segmented into brain regions (Allen Reference Atlas, ARA). All cortical cells in our analyses were recorded from electrode sites verified to be in a motor cortical region. All cells recorded from electrodes located at the pial depth of 1.75 mm or higher (estimated by the manipulator) were assigned a motor cortical region. This depth of 1.75 mm agreed with our physiological estimation of the cortical border (Fig. S2), thus, we considered 1.75 mm as the putative cortical border.

## DATA ANALYSIS METHODS

### Neural data analysis

Single unit data analyses and statistical tests were performed using custom-written codes in Matlab. Spikes of isolated single units in M1 and striatal areas were counted within 1-ms bins to generate the trial-by-bin spike count matrix per unit aligned to reach start or reward delivery. The trial-averaged firing rates were calculated within 50-ms bins and z-score normalized using the mean and standard deviation of its baseline (a 2500-ms period before reach start) firing rate.

### Dimensionality reduction (PCA)

To find the direction along which the neural population activity most covaried during task performance and extract low dimensional neural population trajectories along these directions, PCA was performed on a data matrix D of size (b×t, n), where b and t are the number of 50-ms time bins and the number of trials, respectively, n is the number of neurons. The trial-by-trial binned spike counts are square-root transformed to construct D. Applying PCA to D obtains X and W such that X = DW, where X is the projection of the data onto the principal components (PCs), which are orthonormal columns comprising W that contain the weights from neurons to PCs. To reveal the time-evolving patterns of population activity, D_t,b_ were projected onto top three PCs, trial-averaged and strung together across time to generate neural population trajectories on each PC dimension versus time (Figs. S3-5).

### Linear (consensus) decoders and targeted dimensionality reduction

To assess the contribution of distinct neural populations to forelimb movement, we used a linear decoder to estimate the joystick movement based on the neural activity. The decoded estimates were then compared with held out observed joystick trajectories to assess decoder performance. The decoder defines linear mapping (*W_decode_*) of dimension (N=number of units x D=joystick x,y position) between the neural population activity (F) of dimension (N=number of units x T=time points) and (K) the two dimensional position of the joystick of dimension (D=joystick x,y position x T=time points):

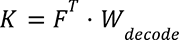

where *F* is the data matrix comprising the population vector of z-scored smooth spike rates (gaussian kernel with *σ* = 24ms; [performance was stable for a broad range of parameters tested]). To solve for a consensus decoder, inspired by general committee machine approaches in machine learning, we solved for the optimal decode vector for batches (data shown used 50 batches of 50 movements each, but good performance was observed from a broad range of parameters settings) in a permuted order and concatenated into a data matrix. For each batch an optimal decoder (minimized mean squared error) was obtained by multiplying the pseudoinverse of the neural data matrix by the movement data matrix. A consensus decoder was obtained by using the mean of the 50 batch decoders (median or centroid can also be used with good performance). To assess performance the decoder was assessed on a small number of held out movements (∼10% of session data). To assess the partial contribution to decoding of specific cell-types or anatomical depth bins we computed the correlation between predicted and actual output using only the weights from the units of interest. These partial correlations were then normalized to the total performance of the decoder for a given session (Fig. 2).

Targeted dimensionality reduction, inspired by previous work (*85*), was approached in a similar manner to the consensus process described above. Previous work calculated beta values of a regression between neural spike counts or averaged activity and scalar task parameters over a task specific time interval. Here, we computed the normalized spike count over the time window that captured the velocity of the outward joystick movement component (see Fig. 1 for velocity profile). We report data from 2 behavioral variables: movement direction (angle of the vector extending from the origin to the position of the peak amplitude displacement) and peak amplitude of the movement. Rather than solving for a single regression coefficient using all data (as before) we again computed coefficients for 50 batches of permuted trials and then took the consensus value. We found this to yield far superior performance to a single regression. Again building upon previous work (*54*), we then sought to identify two orthonormal dimensions of population activity that best captured amplitude (AMP) and direction (DIR) encoding using the Gram-Schmidt method to orthogonalize the consensus decoder dimensions. Again as per previous work (*54*), to calculate tuning along these encoding dimensions we binned movement trajectory data into quintiles for amplitude and direction of movement. For each quintile we computed the mean of behavioral data and the mean of the neural activity weighted by its coefficient for the AMP and DIR dimension. The slope of these 5 points was used to calculate AMP and DIR “tuning”, respectively.

### Naive Bayes classifier

To assess how informative distinct neural populations are of the executed movement amplitude, we used a Poisson naive Bayes classifier to decode which movement amplitude tertile (*C_k_*, *k*=1,2,3) a given trial is sampled from. For each of 1000 iterations, data from each subpopulation (e.g. PT^+^, IT^put^, Striatum etc.) resampled to match the number of neurons per subpopulation are randomly split into 10 folds of trials. A Poisson likelihood function is given by the following:

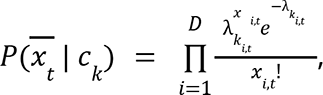

where 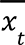 is a population vector of spike counts at *t^th^* time bin of a trial, *k* indicates a movement amplitude tertile. *i* indicates neuron label, 1 to *D*. 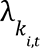 is the parameter for the Poisson distribution estimated using the 9 of the 10 folds by the following:

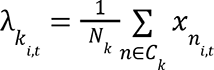

The posterior probability of a movement amplitude tertile given the spike count vector is provided by Bayes’ theore<u>m</u> as follows:

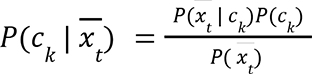

An estimated movement amplitude tertile is assigned to a given trial as follows:

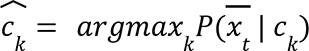

The result of naive Bayes classifier analysis is quantified as the percentage of correctly estimated test folds.

## ACKNOWLEDGEMENTS

The authors thank Brett Mensh, Luke Coddington, and Jason Keller for comments on earlier drafts of the manuscript. The authors thank several members of Janelia Research Campus for assistance with experimental hardware and software. For Neuropixels recordings Brian Barbarits, Jennifer Colonell, Tim Harris, James Jun, Bill Karsh, Wei-Lung Sun, and Eric Yttri provided critical assistance in developing the recording system and associated hardware. Dohoung Kim and Dave (Liu) Liu provided assistance for preparation and analysis of light sheet microscopy data from cleared mouse brains. For imaging experiments Dan Flickinger provided key assistance in the design of the microscope. Janelia Viral Tools and Vivarium provided critical support for these experiments. A.W.H. is a Group Leader and J.T.D. is a Senior Group Leader at Janelia Research Campus of the Howard Hughes Medical Institute (HHMI). This work was supported by funding from HHMI. Competing Interests: The authors declare that they have no competing interests.

## AUTHOR CONTRIBUTIONS

Conception and design of experiments J.W.P., J.P., A.W.H., & J.T.D. Analysis of electrophysiological and behavioral data: J.P. & J.T.D. Data collection for joystick behavioral experiments & optogenetic perturbations: J.P., J.W.P., K.A.M.; Data collection for reach to grasp experiments: J.Z.G.; Data collection for electrophysiological experiments: J.P., J.Z.G. Data collection and analysis for imaging experiments: J.W.P. The first draft of the paper was written by J.P. & J.T.D.; Final version was produced with input from all authors.

## DATA AVAILABILITY

The data used in this manuscript will be made available at <u>janelia.figshare.com</u> (https://janelia.figshare.com/account/home#/projects/125002) upon publication. All data analysis was developed using Matlab (versions 2018b+) or Python (2+). No custom packaged software was produced for these analyses, but useful analysis scripts will be available in FigShare project and/or at our lab repository https://github.com/DudLab and/or at http://dudmanlab.org.

## Supplemental video

**Supplementary video 1. Example trials with/without closed-loop inactivation of MCtx^FL^ neural activity by activation of inhibitory neurons in VGAT-ChR2 mice.** (Left) A representative trial with closed-loop inactivation of MCtx^FL^. A filled circle in the upper left corner indicates frames with laser on. Numbers in the bottom left corner indicate time relative to the laser onset triggered by slight movement of the animal. Video replay is at x0.2 speed.

(Right) A representative trial without closed-loop inactivation. An empty circle in the upper left corner indicates frames for which the laser would have been delivered. Numbers in the bottom left corner indicate time relative to the pseudo laser onset.

Video 1 can be found online at: https://www.dropbox.com/s/vy6dzrqwm4w1eix/Trial%2354_139_stim-Pstim%20%28Converted%29.mov?dl=0

**Supplementary video 2. Inactivation of IT neurons triggered by the hand lift discontinues an ongoing reaching.**

The video provides dual views on a mouse performing a reach-to-grasp task. To highlight the necessity of continuous IT neuronal dynamics for a successful forelimb reach, we automatically triggered a laser at the detection of a voluntary hand lift, which inactivated IT neurons using the inhibitory opsin GtACR2. The laser onset is signaled by a LED light (invisible to the mouse) in the video. IT inactivation discontinued the ongoing reaching and blocked subsequent reaching during inactivation which lasted for 2 s.

Video 2 can be found online at: https://www.dropbox.com/s/1sayuypoujj6ir1/IT_liftTriggeredSilencing_20200125_v007.mp4?dl=0

**Supplementary video 3. Inactivation of PT neurons of MCtx^FL^ has little impact on forelimb movement in a reach-to-grasp task.**

The current trial begins with a cue signaling placement of the pellet in the target position which coincided with the onset of a laser that inactivated pons-projecting PT neurons using the inhibitory opsin GtACR2. The cue/laser onset is signaled by a LED light. PT inactivation had little impact on reaching.

Video 3 can be found online at: https://www.dropbox.com/s/rgpppdmlid12qea/PT_silencing_20200426_v120.mp4?dl=0

---

**Fig. S1.**
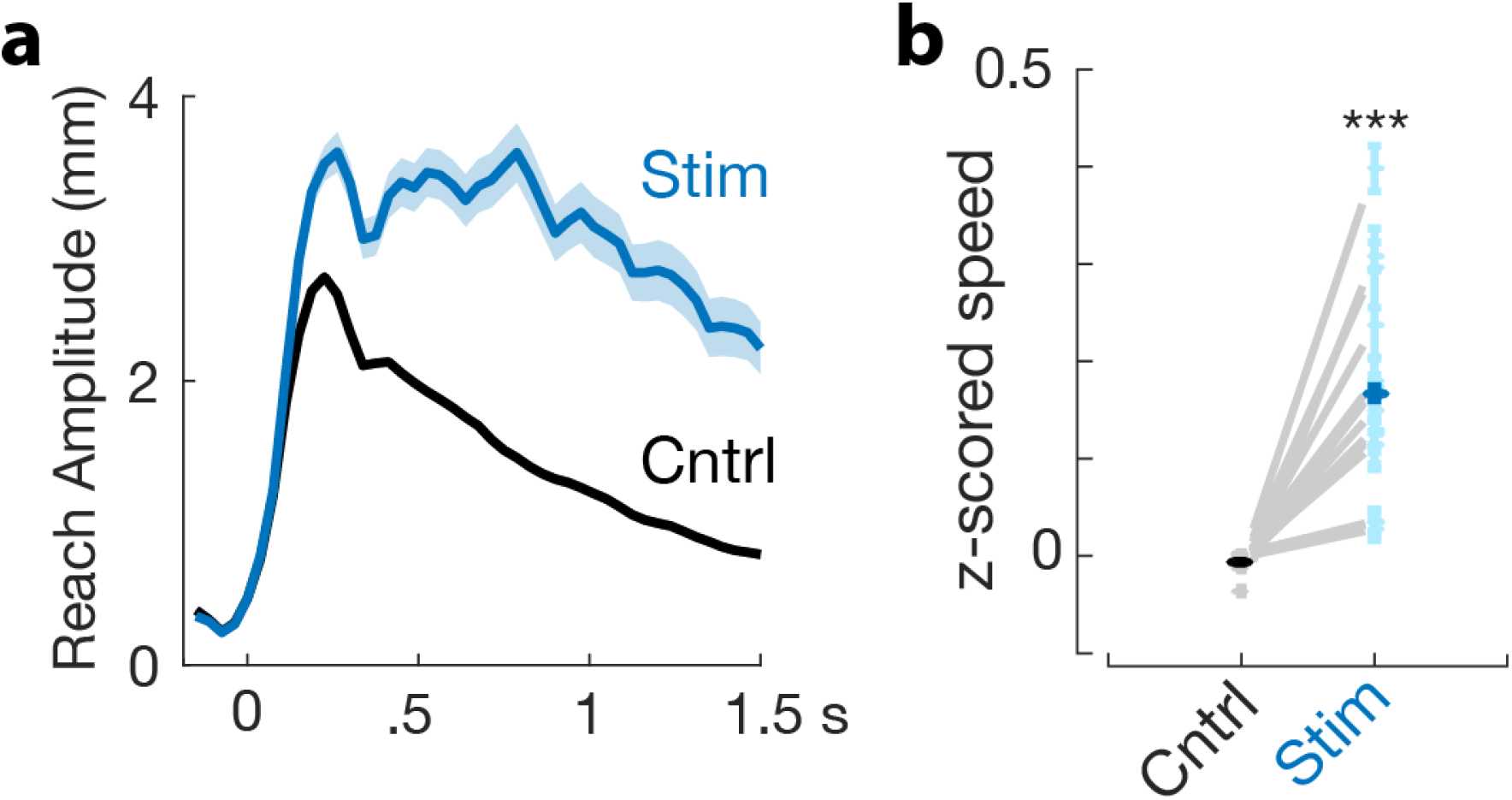
Stimulation of layer 5 output neurons in MCtx^FL^ invigorates forelimb movement. Closed loop stimulation of the majority of descending layer 5 output neurons in MCtx^FL^ labelled using the Rbp4-cre line (*48, 67*) crossed to Ai32(*76*) produced increases in the amplitude (*top*) and speed (*bottom*) of forelimb movement.

**Fig. S2.**
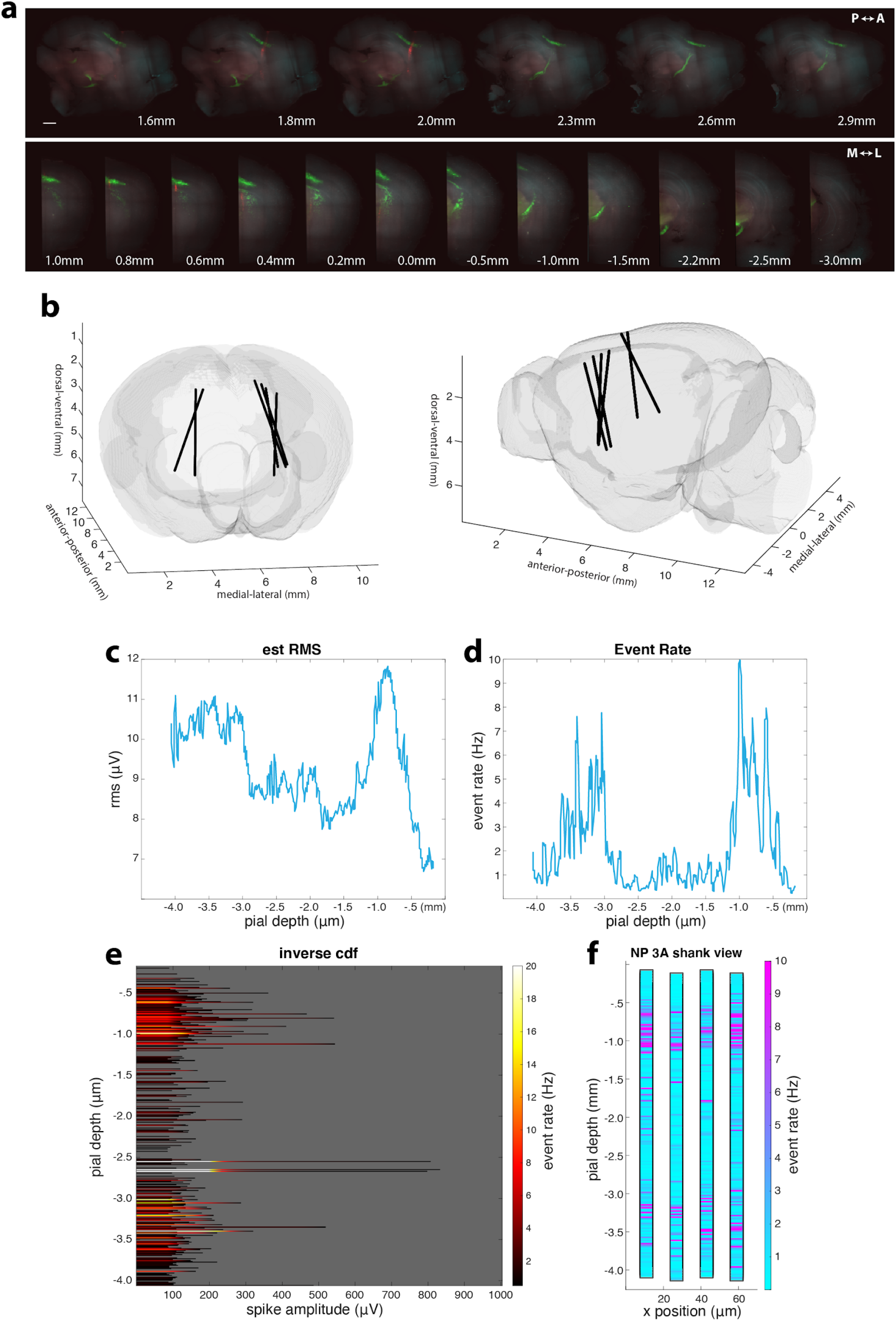
Histological verification of Neuropixels recordings. **a,** A sagittal (top) and coronal (bottom) view of a cleared mouse hemi brain imaged with light sheet microscopy. Green fluorescence indicates labeling of the deep layer 5 PT neurons and their projections to downstream areas such as striatum, superior colliculus and pons. Red fluorescence indicates probe tracks. Numbers in the top and bottom rows indicate medial-lateral and anterior-posterior coordinates relative to bregma, respectively. The length of the white scale bar = 1mm. **b**, Three dimensional rendering of probe tracks with the Allen Anatomical Template (AAT). Imaged 3D brain volumes were aligned to AAT, and each electrode on Neuropixels probe was assigned to a brain region using the probe geometry and Allen Reference Atlas (Methods). All cells recorded from electrodes located at the pial depth of 1.75 mm or upper (estimated by the manipulator) were assigned a motor cortical region. **c**, Estimated root mean square voltage (RMS) is plotted versus the pial depth. Note the trough indicating dearth of neural activity around the depth of 1.75 mm, which is consistent with the histologically estimated cortical border. **d**, Mean event rate is plotted versus the pial depth. An event is defined as voltage crossing (e.g. spikes) a threshold (80 µ*V*). Note the elevated event rates of the cortical depths. **e**, Threshold-crossing events are binned and counted based on the absolute raw amplitude for each pial depth. **f**, Mean event rates are plotted for each electrode site tiling the Neuropixel probe. Note the higher event rates within the cortical range as well as the dearth of events around the histologically estimated cortical border (1.75 mm). RMS and event rates were measured using codes written by Jennifer Colonell (https://github.com/jenniferColonell/Neuropixels_evaluation_tools).

**Fig. S3.**
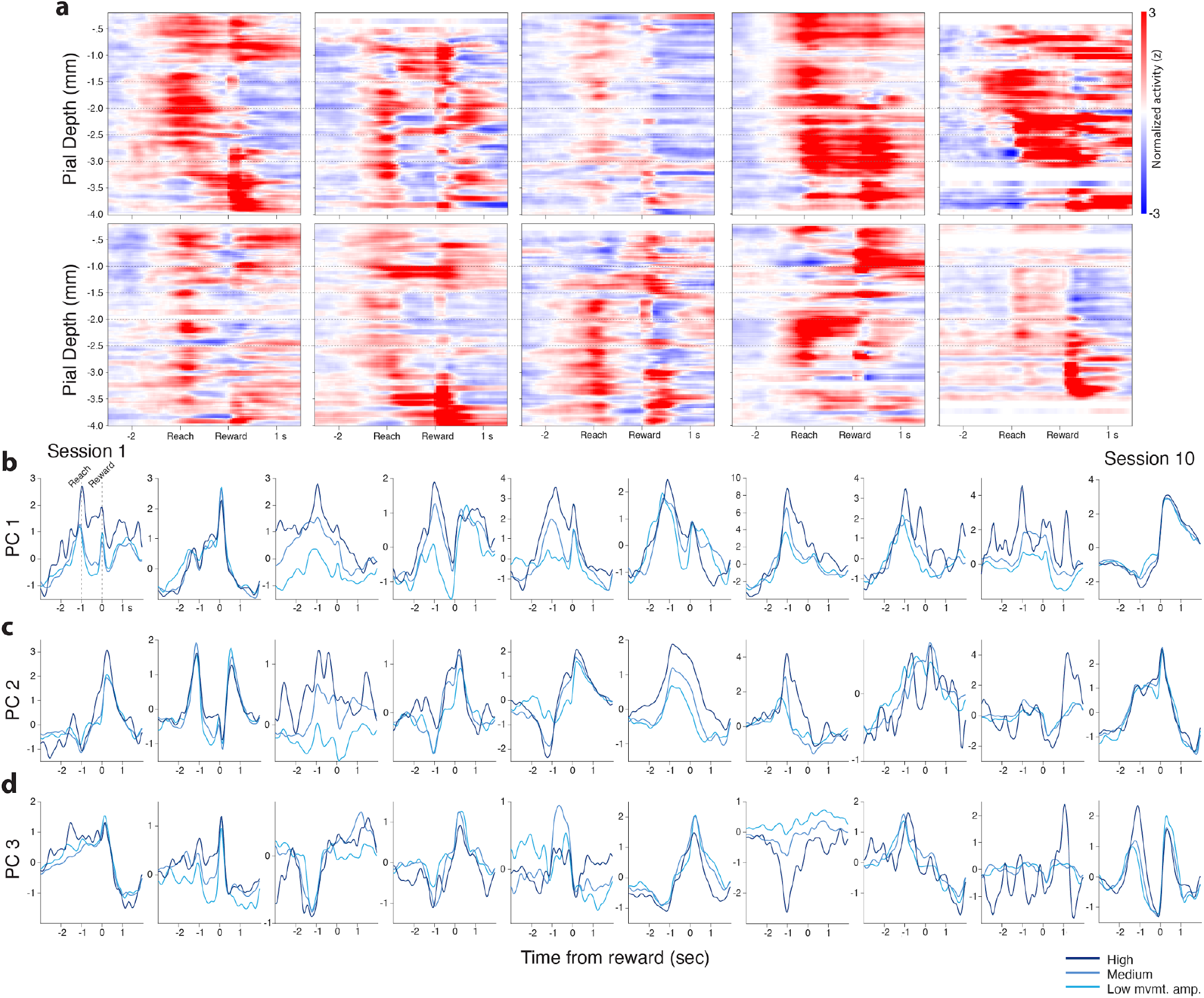
Summary of cortical and striatal neural population activity during task performance. **a**, Mean z-score normalized activity aligned to reach threshold crossing and reward delivery (x axis) is plotted per inferred depth (y axis) relative to the pial surface for all individual recording sessions (N=10). **b-d**, MCtx^FL^ neural population activity of movement amplitude tertiles are projected onto the top three principal components. Neural trajectories indicate that at least partially separable populations of neurons are active during forelimb movements scaling with reach amplitude. Other populations appear to be active during reward collection.

**Fig. S4.**
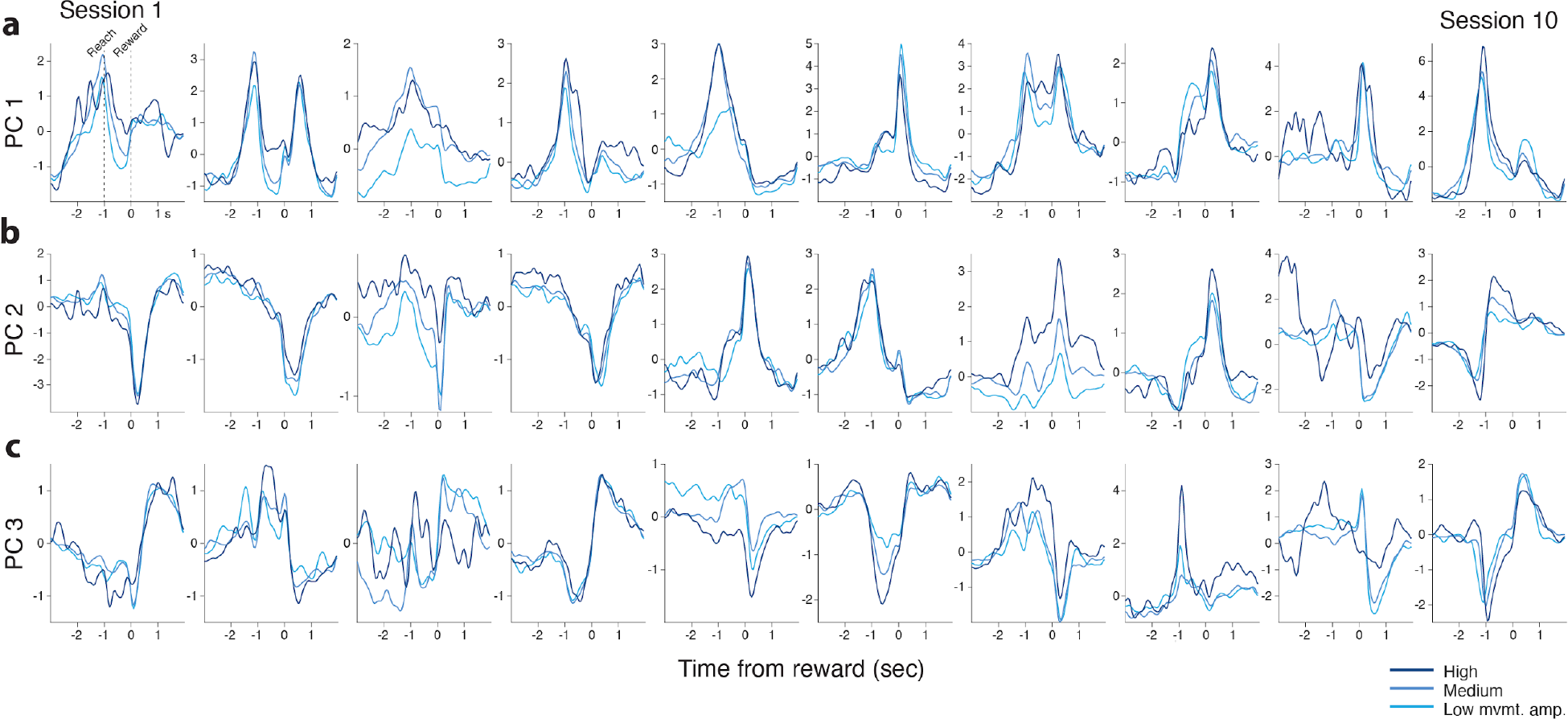
Projection of striatal neural population activity onto top three PCs. **a-c**, Striatal neural population activity of movement amplitude tertiles are projected onto the top three principal components. Similar to the cortical data in Fig. S4, neural trajectories indicate that at least partially separable populations of neurons are active during forelimb movements or reward collection.

**Fig. S5.**
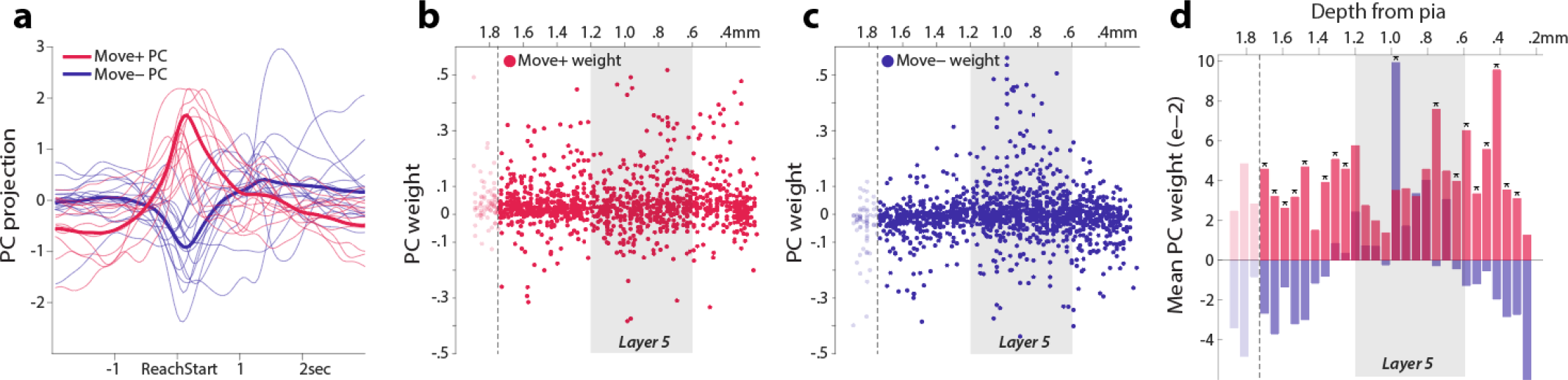
Laminar inhomogeneity of movement-related activity. **a**, MCtx^FL^ neural activity projected onto the top three PCs (Methods). PCs were classified as ‘Move+’, if the neural trajectory on each PC dimension (PC score) peaked before +500 ms relative to the reach start. Other PCs that peaked afterwards with negative modulation during reach were classified as ‘Move-’. **b**, Weights of move+ PCs are plotted as a function of recording depth across all cortical neurons (solid circles). A dotted line indicates the histologically-verified cortical border (1.75 mm, see Methods and fig. S3). **c**, Weights of move-PCs are plotted as a function of recording depth. **d**, Weights of move+ and move-PCs significantly differ across cortical depths (Two-way ANOVA, F_1,29_ = 2.15, p = 4.0×10^-4^, bin-by-bin test, p < 0.05).

**Fig. S6.**
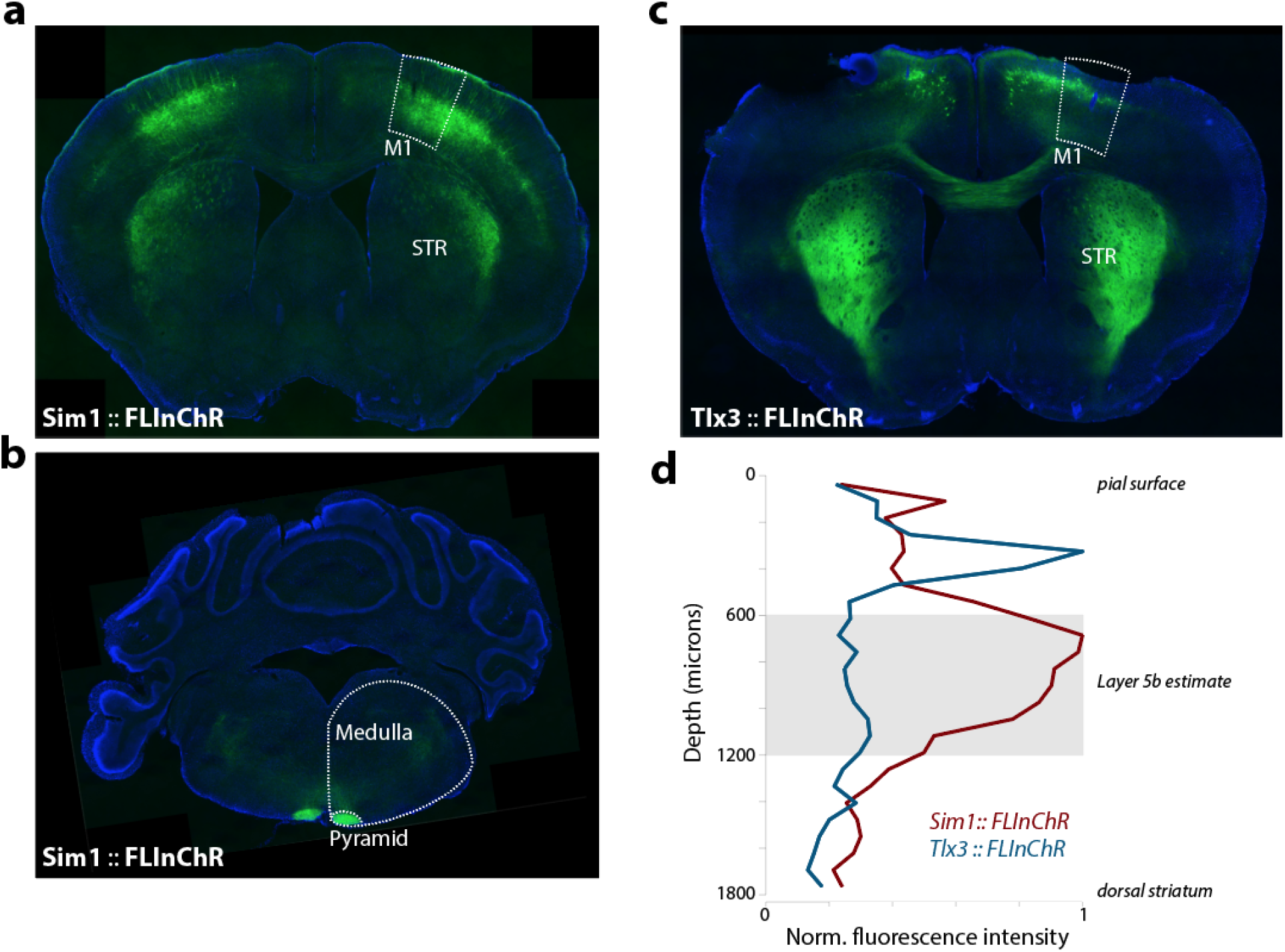
Histological verification of PT and IT neuronal labelling. **a**, Expression of fluorescently-labelled FLInChR in the deep layer 5 PT neurons of Sim1-cre mice. **b**, Fluorescent labelling of the medullary pyramid in PT mice. **c**, Expression of FLInChR in the layer 5 IT neurons of Tlx3-cre mice. **d**, To compare the laminar distribution of PT and IT neurons in the motor cortex, the intensity of fluorescence was measured along a line oriented from the pial surface through M1 to the dorsal surface of STR. This distance was normalized to 1800 microns which is our estimate from the allen mouse brain atlas. The fluorescence intensity data were binned into 250 evenly spaced bins and averaged within each bin. The binned data were normalized to max intensity for each cell type.

**Fig. S7.**
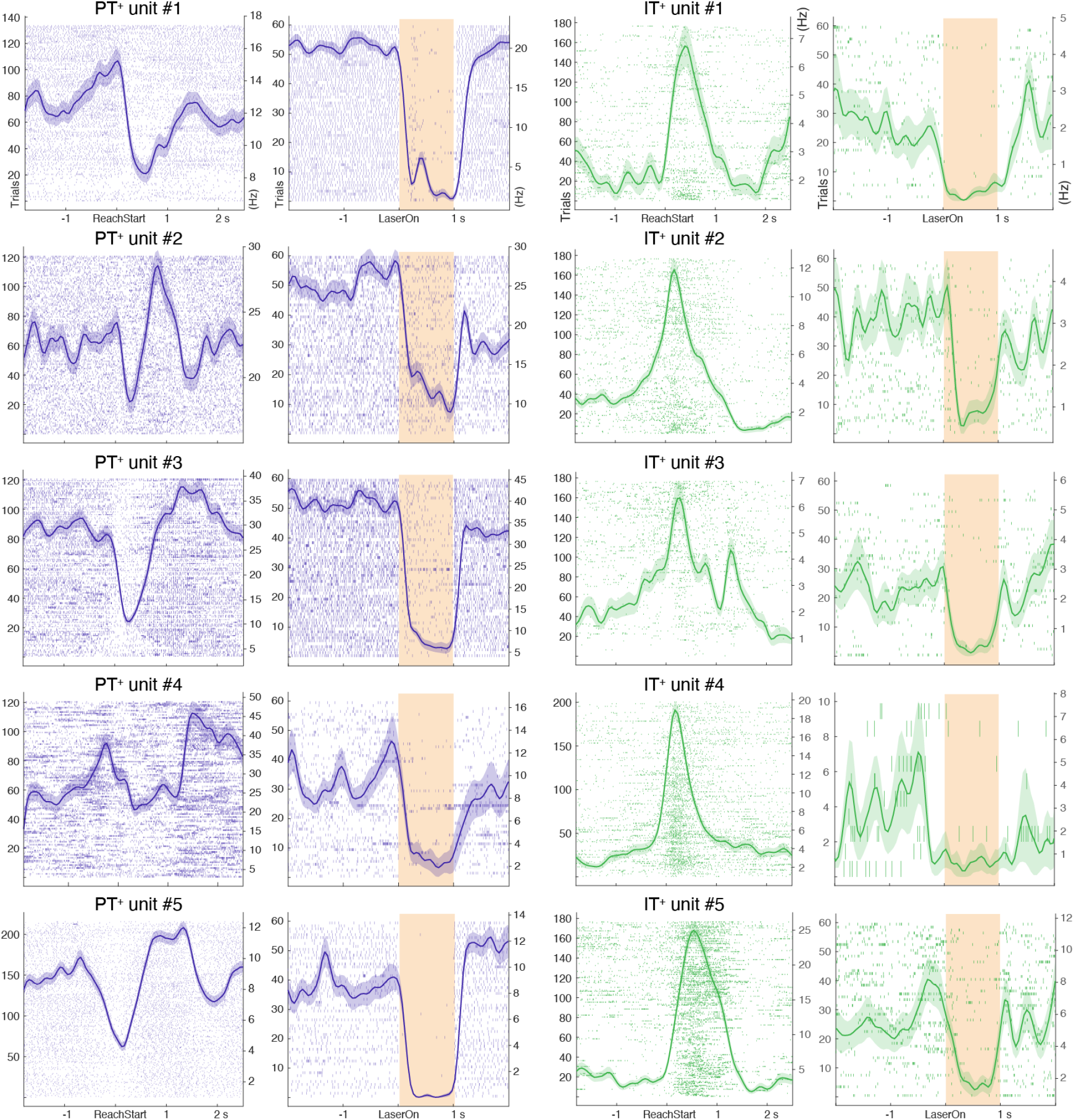
PT^+^ and IT^+^ neuronal activity during task performance and optotagging. Blue-colored rasters in the left column illustrate trial by trial individual neuronal responses during task performance (left panel of each pair, aligned to reach start) with significant inhibitory responses during optotagging (right panel of each pair, aligned to laser onset) in Sim1-Cre (KJ18Gsat) mice injected with rAAV2-retro-CAG-Flex-FLInChR-mVenus to the pons. Each row represents each trial. The mean±SEM spike rate (Hz) is superimposed. Numbers on the left and right ordinates of each plot indicate the number of trials and firing rate in Hz, respectively. Green-colored rasters in the right column illustrate trial by trial IT^+^ individual neuronal responses during task performance (left panels) and optotagging (right panels) in Tlx3-Cre (PL56Gsat) mice.

**Fig. S8.**
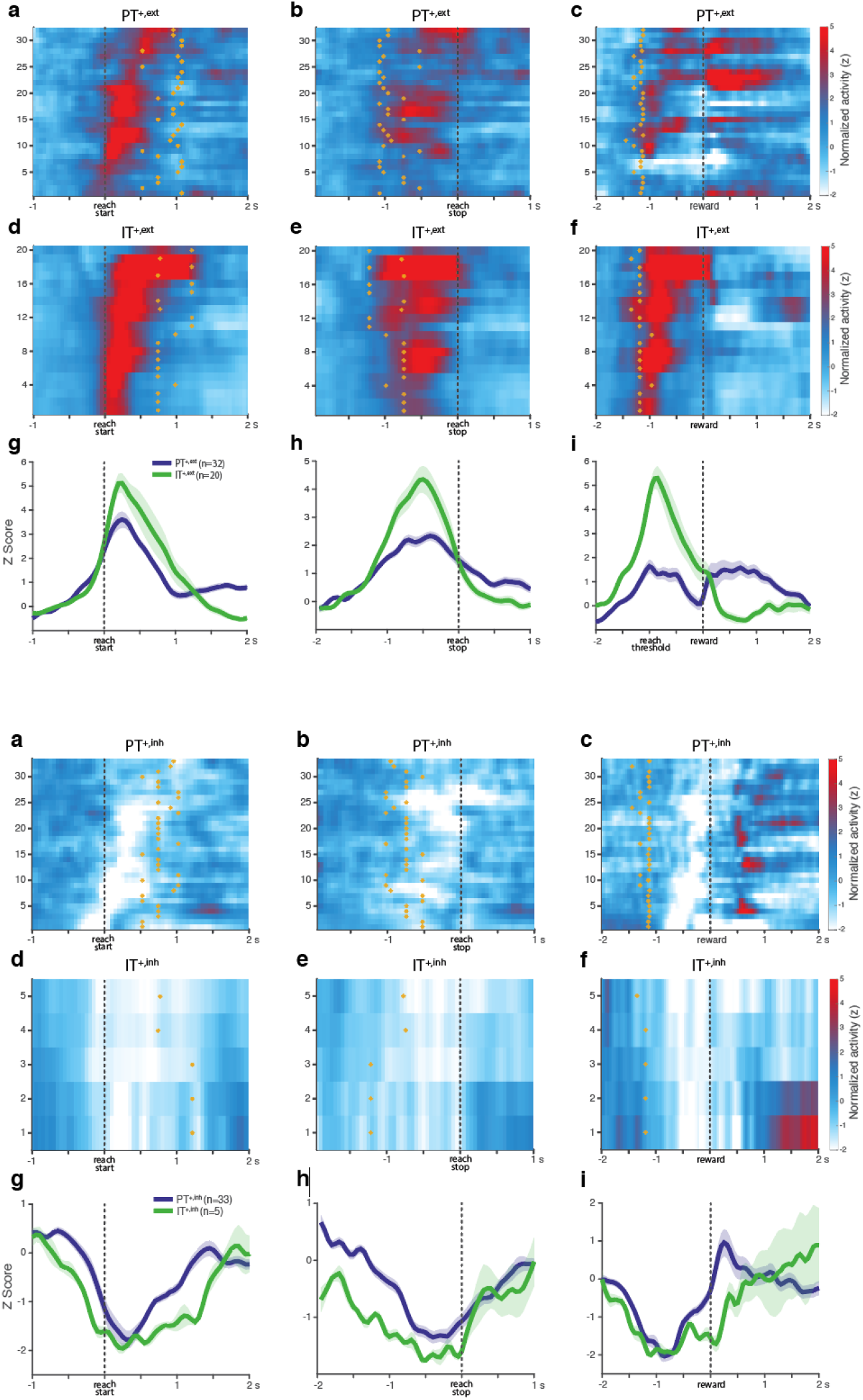
Activity of PT^+^ and IT^+^ neurons with a significant positive modulation of activity during movement aligned to different movement phases. *Upper:* **a,** The mean z-score normalized activity of PT^+,ext^ neurons (n=32 of 111 PT^+^) aligned to reach start. To indicate the mean duration of movement an asterisk was superimposed on each row at the time bin where reach stop occurred. **b,** Activity of the same PT^+,ext^ neurons but aligned to reach stop. Asterisks indicate the mean time of reach start. Neurons were identically sorted along the y-axis across panels (a-c). **c,** Activity of the same PT^+,ext^ neurons but aligned to reach threshold crossing and reward delivery. Note that there was a constant 1-sec delay between the reach threshold crossing and the reward delivery. Asterisks indicate the time of reach start. **d,** Activity of IT^+,ext^ neurons (n=20 of 30 IT^+^) aligned to reach start. **e,** Activity of IT^+,ext^ neurons alignmed to reach stop. **f,** Activity of IT^+,ext^ neurons alignmed to reach threshold crossing and reward delivery. **g-i,** For comparison, the mean±SEM normalized activities of PT^+,ext^ and IT^+,ext^ neurons were plotted with alignment to different movement phases. *Lower:* same as Upper but for negative modulation around movement.

**Fig. S9.**
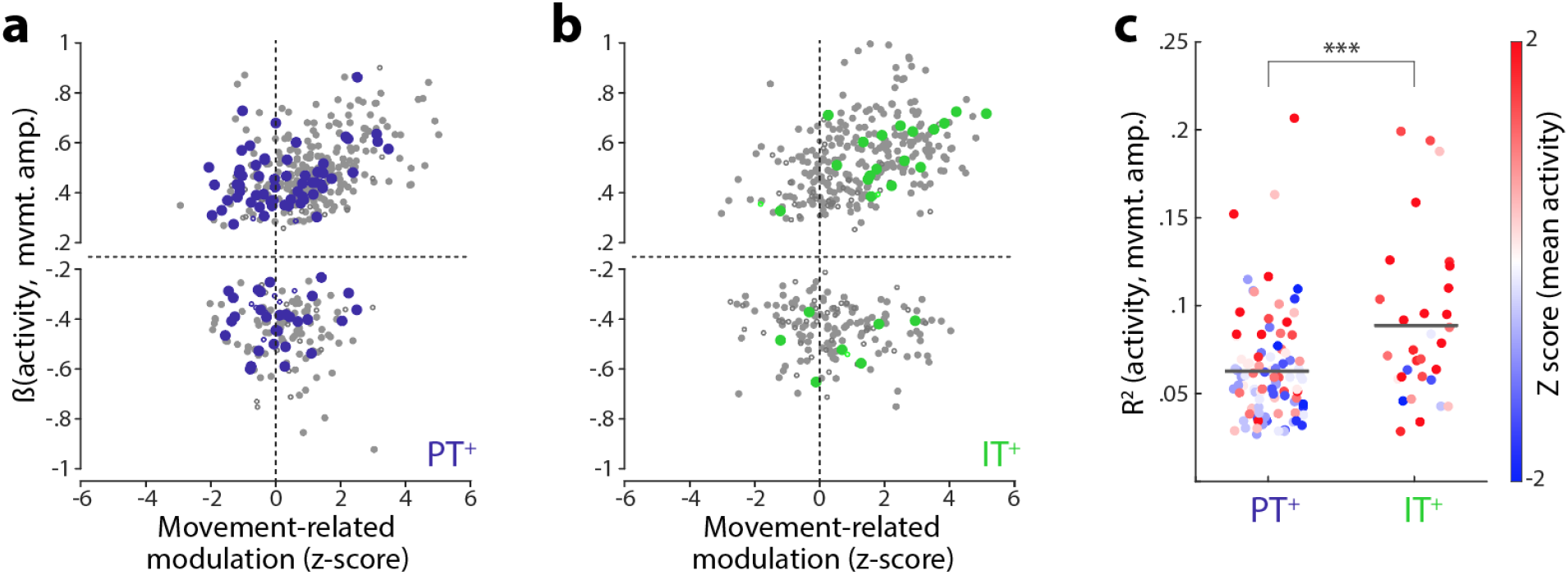
Cell-type specific individual neuronal correlation with movement amplitude. **a**, Individual PT^+^ units are plotted based on their normalized movement-timed activity along X axis and their activity modulation as a function of movement amplitude along Y axis (regression coefficients, β). For instance, a unit in the first quadrant is the one that increased its firing rate during movement with a positive correlation with the movement amplitude. Blue-filled circles represent PT^+^ units with a significant regression coefficient (*t* test, α=0.05). Gray circles represent the rest of MCtx^FL^ units that were not tagged. Fisher’s exact test against a uniform distribution of PT^+^ units across quadrants, p=0.84). **b**, The vast majority of the individual IT^+^ units located in the first quadrant (Fisher’s exact test, p=0.0056). Green-filled circles represent IT^+^ units with a significant regression coefficient. Gray circles represent the rest of MCtx^FL^ units that were not tagged. **c**, Squared Pearson correlation between normalized firing rate and reach amplitude are plotted for individual PT^+^ and IT^+^ units color-coded by their mean movement-timed activity.

**Fig. S10.**
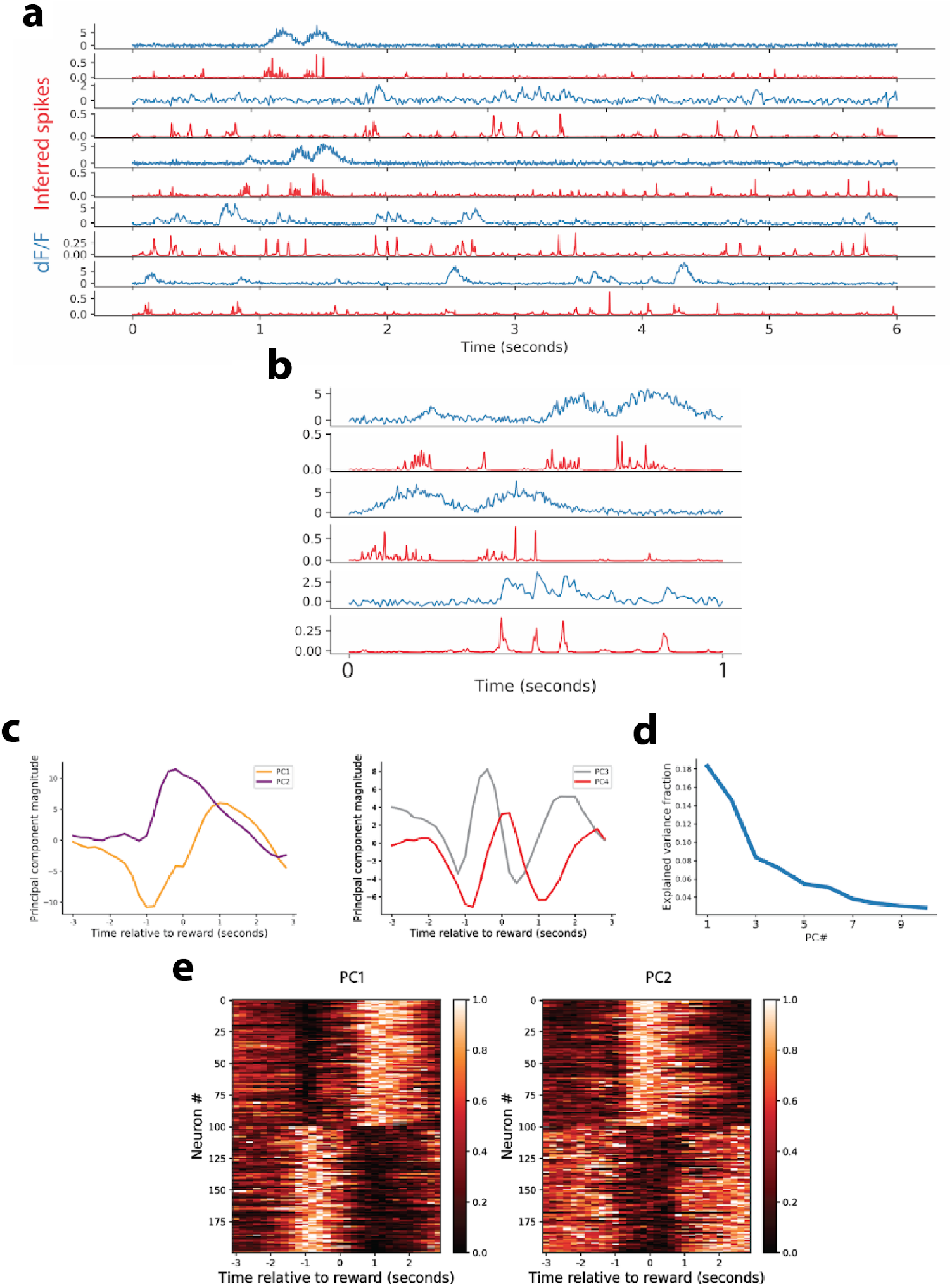
Spike deconvolution and PCA for calcium imaging data. **a**, Illustration of spike deconvolution. Panel shows 5 example regions of interest, each of two rows. Top row for each unit (blue) shows the dF/F trace, with the row beneath (red) showing the inferred spike activity metric. **b**, a zoomed-in portion for three units from **a**. **c**, Structure of first four principal components of neural activity across all units in the dataset, aligned to reward. **d**, The fraction of explained variance for the top 10 principal components in the dataset. **e**, *left*, The normalized inferred spike rates of representative units with most positive (top 100 rows) and negative (bottom 100 rows) weights for PC1 are plotted. Format same as Fig. 3e. *right*, The normalized inferred spike rates of representative units with positive (top 100 rows) and negative (bottom 100 rows) weights for PC2 are plotted.

**Fig. S11.**
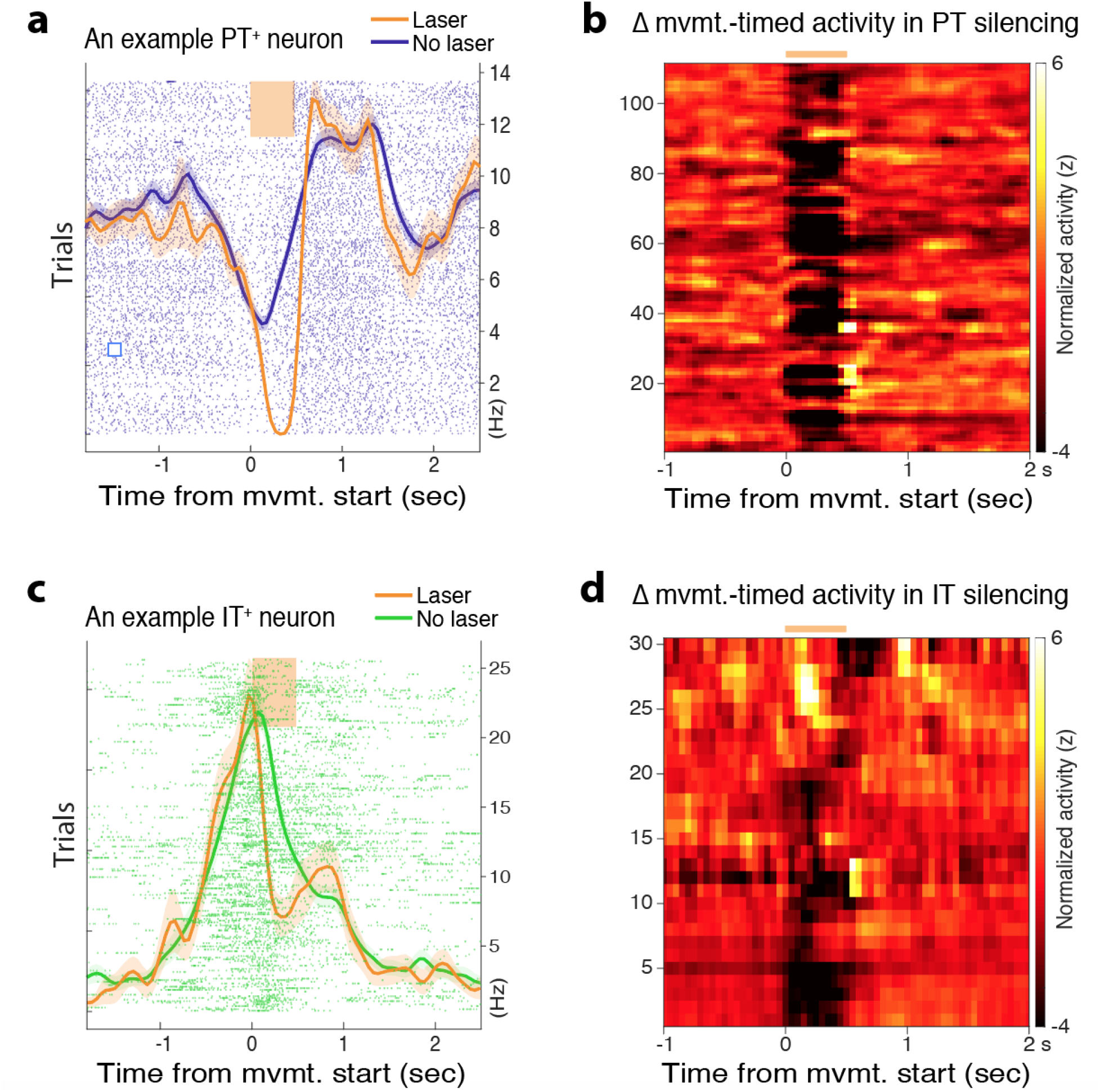
Robust PT^+^ and IT^+^ neuron inactivation during task performance. **a**, An example PT^+^ neuron displays a robust inactivation by the laser triggered at the earliest detection of reach in randomly selected trials (top raster rows: laser trials; bottom raster rows: control trials). The mean±SEM spike density (Hz) functions are superimposed for laser and control trials. **b**, Change of all individual PT^+^ neuronal activity by opto-silencing aligned to the movement onset. **c**, An example IT^+^ neuron displays a robust inactivation by the laser. **d**, Change of all individual IT^+^ neuronal activity by opto-silencing.

**Fig. S12.**
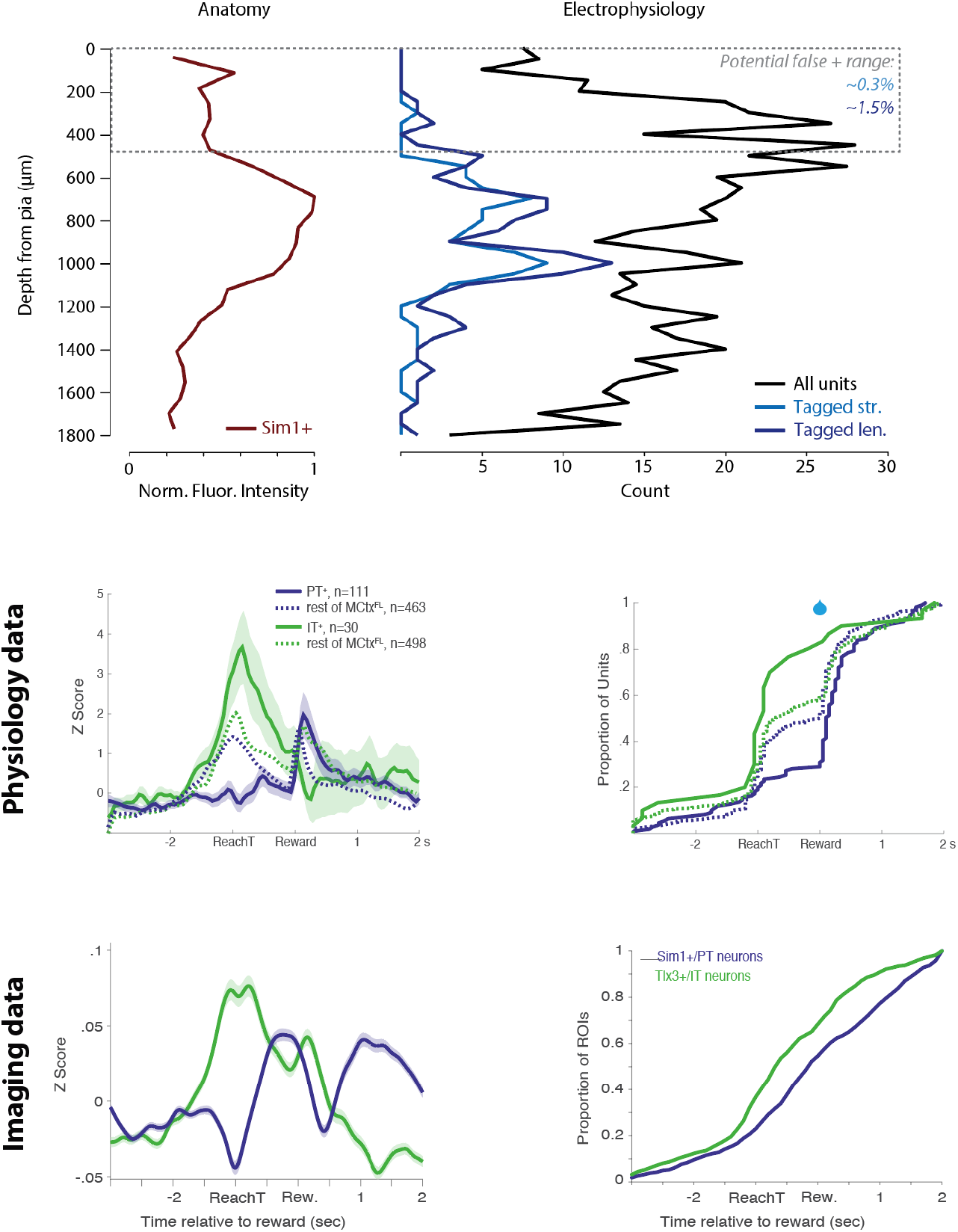
Comparing anatomical distribution of cells with putative identified cell types from optotagging. These datasets are almost certainly still dominated by false negatives - individual units that are representative of a given cell type, but fail to meet inclusion criteria from optotagging. However, there is also the possibility that a given unit shows strong modulation during the tagging procedure (inhibition to illumination consistent with expression of FLInChR). Our most stringent (‘str.’) inclusion criteria include mean suppression of firing below −0.3 z score units (z score defined over entire recording duration) sustained for the entire duration of illumination (based upon evidence that FLInChR can produce non-desensitizing inhibition for >500 ms (*53*)). Our more lenient criterion did not require sustained inhibition for the entire illumination duration (‘len.’). All results based upon tagged units in the paper are confirmed to be consistent over a range of stringencies. We consider the false positive rate at this stringency which is used for results from key behavior-physiology sessions used for comparison of cell type differences in encoding in Figure 4 in particular. Here we align estimates of PT (Sim1+) neuron density from fluorescent intensity at left (see Fig. S6) compared to the count of PT+ neurons as a function of depth and the count of all cortical units recorded in the same sessions. ∼6% of all units recorded were identified as tagged in these datasets. The true number of Sim1+ neurons out of the recordable population is not known and may exceed 6%. One estimate of the false positive rate lower bound is to consider a range of anatomical depths at which no Sim1+ somata are thought to be located (<500 microns in depth) indicated by an inflection in fluorescence density and consistent with prior labeling work (*2*). In our imaging experiments we typically imaged at ∼400 µm depth (Fig. 5) and saw little to no somata at this depth for Sim1+ neurons. In this region we recorded 330 single units of which 1 was deemed tagged (str.) or 5 tagged (len.) giving a putative lower bound on false positive rates at 0.3%-1.5%. These estimates amount to removing one or a couple units from distributions on tagged units and there is little impairment to conclusions from such changes. We note that localization of a unit on a Neuropixels probe is not necessarily the somata (proximal dendrite has large extracellular currents during spiking) and thus we do not know that the true positives are 0. Lastly, we also show a direct comparison between tagged unit average properties and imaged populations with a nominal 0% false positive rate. These are replotted from Figure 3 and 5.

## Notes

https://janelia.figshare.com/account/home#/projects/125002

### Competing Interest Statement

The authors have declared no competing interest.

### Summary of Updates

Several figures have been edited with new analyses, the addition of some a new dataset, and a modest rearrangement of figure organization

